# Hippocampus maintains a flexible and coherent map under reward flavor-landmark cue conflict

**DOI:** 10.1101/2021.09.15.460484

**Authors:** Indrajith R. Nair, Dipanjan Roy

## Abstract

Animals predominantly use salient visual cues (landmarks) for efficient navigation over other sensory modalities. When the relative position of the visual cues is altered, the hippocampal population exhibits heterogeneous responses and constructs context-specific spatial maps. Another critical factor that can strongly modulate spatial representation is the presence of reward. Reward features can drive behavior and are known to bias spatial attention. However, it is unclear whether reward flavors are used for spatial reference in the presence of distal cues and how the hippocampus population dynamics changes when the association between reward flavors and distal cues is altered. We investigated these questions by recording place cells from the CA1 while the rats ran in an environment with the conflicting association between reward flavors and distal cues. We report that the hippocampal place cells coherently and dynamically bind to reward flavors or distal cues across sessions, but not simultaneously suggesting the use of a single spatial map. We found that place cells maintained their spatial offset in the cue conflict conditions, thus showing a robust spatial coupling featuring an attractor-like property in the CA1. When the textures were added on the track, the coherency of the CA1 is degraded, as the hippocampus showed a heterogeneous response and weak spatial coupling of co-recorded cells suggesting a break away from the attractor network. These results indicate that reward flavors alone may be used for spatial reference but may not cause sufficient input difference to create context-specific spatial maps in the CA1.

## Introduction

Animals use multimodal information to encode the space (Jacobs & Schenk, 2003; Cushman *et al*., 2013), and the hippocampal system receives information from various sensory modalities (Ranck, 1973). The hippocampal encoding of space is thus multifactorial, where place cells use a repertoire of salient external sensory cues and internal network dynamics (attractor states) to organize the knowledge of any change in the environment (O’keefe and Speakman, 1987; Shapiro *et al*., 1997; Samsonovich & McNaughton, 1997; Tsodyks, 1999; Wills *et al*., 2005; Knierim & Neunuebel, 2016). When animals encounter different environments, the hippocampus responds to it with a sparse, orthogonal representation, where different combinations of place cells are recruited to represent each environment (Alme *et al*., 2014). Also, individual neurons in the CA1 respond differently to a change in the spatial association between external stimuli showing a hierarchical representation of cues (Shapiro *et al*., 1997; Knierim,2002; Lee *et al*., 2004; Yoganarasimha *et al*., 2006). Place cell orientation is acutely controlled by visual cues, where the place fields rotate with the rotation of the landmarks (O’keefe & Conway, 1978; Muller & Kubie, 1987). In the absence of distal cues, place cells prefer local visual cues for context-specific spatial representation (Draht *et al*., 2017). Also, only in the absence of light, odor cues orient the place fields and exhibited remapping when odor cue orders were scrambled (Save *et al*., 2000; Zhang & Manahan-Vaughan, 2015; Lebedev & Ossadtchi, 2018). These studies indicate the preference of the hippocampal system in using visual cues over other sensory modalities as a reference frame to navigate the environment seamlessly (Cushman *et al*., 2013).

In addition, hippocampal place cells can also efficiently encode non-spatial information such as visual cues (Fried *et al*., 1997), odor (Wood *et al*., 1999), taste (Herzog et al., 2019), textures (Shapiro *et al*., 1997; Wang *et al*., 2020), and time (Kraus *et al*., 2013). The hippocampal neurons also respond to ethologically relevant stimuli like reward and goal position to guide motivated behavior. Reward-related modifications are also shown in the Entorhinal Cortex(EC), which provides cortical input to the hippocampus (Boccara *et al.,* 2019; Butler *et al.,* 2019).

Many studies have suggested the over-representation of goal locations and increased population firing at the goal ((Lee *et al*., 2006; Hok *et al*., 2007; Hayashi *et al*., 2016; Tryon *et al*., 2017). Other studies have also shown increased hippocampal activity while approaching the goal (Eichenbaum *et al*., 1987; Breese *et al*., 1989). Recently, Gauthier and Tank showed that a small population of neurons in the hippocampus (reward cells) shifted their firing field with the change of location of the reward but resisted remapping during any change in the context (Gauthier &Tank, 2018). In addition, CA1 pyramidal cells carry both the reward and spatial information to various degrees (Xiao *et al*., 2020). Studies have also shown that the hippocampus transmits contextual information to the mid-brain dopaminergic system to regulate motivational behaviors (Luo *et al*., 2011).

CA1 neurons show the decorrelation of place field activity in the different environments (Alme *et al*., 2014). In addition, CA1 neurons exhibit heterogeneous responses in a dynamically changing environment where a fraction of place cells is oriented to distal or some with local cues ((Knierim, 2002; Lee *et al*., 2004; Yoganarasimha *et al*., 2006). This mixed response, along with the remapping of place fields in different contexts, enables the hippocampus to establish environment-specific spatial maps. However, in most experiments, distal cues are of visual modality, and local cues are usually multimodal, having visual combined with tactile or olfactory features (Shapiro *et al*., 1997; Knierim, 2002). Studies on bats have shown that hippocampal representation showed remapping by shifting the CA1 place fields while switching between two sensory modalities - vision and echolocation (Geva-Sagiv *et al*., 2016).

Many studies suggest that even macrosmatic animals mainly used distal landmarks for spatial and directional reference (Jeffery, 1998). Other sensory stimuli like auditory cues, odor, taste, and tactile information might be only used for spatial reference when the input from the visual modalities was scarce or when the task demanded the usage of such information. However, reward drives and modulates the behavior and may exert an intense evolutionary pressure to associate context and rewarding stimuli for survival. It is also shown that learned association of visual stimuli with the reward is essential in shaping attention (Camara *et al*., 2013; Anderson 2013; Chelazzi *et al*.,2013). The attention to reward also increases the stability of the place fields, modulates place cell discharge (Kentros *et al*., 2004, Moser 2004, Fenton *et al*., 2010) and bias hippocampal replay (Bendor *et al*., 2012; Carey *et al*., 2019). Thus, visual modality and reward features can be considered two strong and sometimes competitive factors in the spatial navigation. Visual cues are predominantly used for spatial reference, whereas reward directs motivated behavior, attention, and decision making. But, it is unclear how hippocampal representation is driven by these two different cue systems when the association between the distal and reward features are altered. In the present study, we systematically investigated the dynamics of hippocampal representation in a dynamically changing reward flavor-distal cue association by recording from the CA1 region of the dorsal hippocampus. We report that CA1 ensembles exhibited a highly coherent yet unstable spatial representation without local cues showing attractor network property. In the presence of local visual cues, the hippocampal neurons show orthogonal activity in a dynamically changing environment, destabilizing the attractor network.

## Materials and Methods

### Subjects

6 Male Long-Evans rats (5-6 months old) were housed separately on reversed light-dark cycle (12:12 h). Food and water were provided ad libitum. All the experiments were conducted on the dark phase of the cycle. All procedures related to surgery, animal care, and euthanasia were approved by the Institutional Animal Ethics Committee (IAEC) of National Brain Research Center at Manesar, Haryana, constituted by the Committee for the Purpose of Control and Supervision of Experiments on Animals (CPCSEA), Government of India and were conducted in accordance with NIH guidelines.

All the surgical procedures were performed under aseptic conditions. At the time of surgery, rats were initially anesthetized with ketamine (60 mg /kg b.w.) and xylazine (8 mg /kg b.w.) and subsequently switched to isoflurane gaseous anesthesia. A custom-built Microdrive containing 9- 20 independently moving tetrodes (1-2 references in each Microdrive) was surgically implanted over the right hemisphere to access dorsal CA1 of the hippocampus (coordinates 3.2-4.8 mm posterior to bregma, 1.2-2.8 mm lateral to midline). To relieve pain, Meloxicam was administered intramuscularly (1 mg/kg b.w.) on the day of surgery. Meloxicam was also provided through the oral route (1 mg/kg b.w.) during the post-surgical recovery period. During the post-surgical care of 7 days, food and water were provided ad libitum. Rats were also provided with different reward flavored pellets. Following the recovery period, tetrodes were lowered into the brain to reach dorsal CA1 of the hippocampus. Rats were trained to run clockwise (CW) on a circular track to receive specific reward flavors at specific spokes on the track. During training and experimental sessions, the rats were maintained to no less than 85% of their body weight.

### Electrophysiological Recording

17 μm VG bonded platinum-iridium wire was used to make tetrodes (California Fine Wire, USA). The tips of individual wires were cleaned to remove debri by passing current (0.2 μA). Then, the tetrode tips were electroplated with the platinum black solution to reduce the impedance to 100-150 k ohm with 0.2 μA current. We used 96 channel data acquisition system to acquire electrophysiological recordings (Digital Lynx 10S, Neuralynx. Inc., USA). The custom-made microdrive was attached to the EIB-27 board. The tetrodes and the references were connected to this EIB board. Headstage preamplifier (HS-27) was connected to the EIB-27 board. HS-27 was used to amplify the brain signals. Using specific tethers, HS-27 was connected via the commutator cables to the data acquisition system. The multi-unit activity was obtained by filtering the brain signals between 600 Hz and 6KHz. Spike waveforms were sampled for 1ms at 32kHz. The spikes waveforms were differentially recorded against a reference electrode present in the corpus-callosum (cell-free layer). Local Field potentials (LFP) were differentially recorded against a ground screw on the skull above the frontal cortex. LFP signals were recorded as continuous signals with a bandwidth between 0.1Hz-1 kHz and continuously sampled at 4 kHz. In this study, only the spike data is used. LFP signals were only used to visually observe ripples and complex spiking patterns while the tetrode tips were reaching the hippocampus, and no analysis was performed on it. The rat position was tracked during experimental sessions using LEDs connected to the headstages, acquired at 25 Hz using a ceiling-mounted camera (CV- S3200, JAI Inc., San Jose, USA).

### Experimental Design

After post-surgical recovery (∼7 days), tetrodes were slowly advanced to reach the CA1 layer of the hippocampus over 12-21 days. The rats were trained for 12-14 days (∼30 min/day) in a behavioral room to run clockwise (CW) on a centrally placed plain circular track (90 cm from the floor) to receive specific flavored pellets as a reward at specific spokes. Whenever the rat tried to turn back and run in a counter-clockwise (CCW) direction, the path was blocked using a cardboard sheet. The circular track (56 cm inner diameter,76 cm outer diameter) had three spokes projecting outwards from the track. The three spokes are separated by 120°. The circular track and the spokes connected to it are black colored. Three distinct flavors of reward pellets– namely banana flavoured, sucrose, and chocolate-flavored pellets are provided at each spoke, respectively (Banana #F07257, Sucrose #F06233, Chocolate #07256, BioServ (Flemington, NJ, USA) Dustless Precision Pellets, 45 mg). Three salient and visually distinct cues (made of cardboard with distinctive pattern and shape) are hung on the black curtain. These three distal cues are aligned in line with the three spokes of the track such that banana flavor spokes are aligned with rectangular stripe card, sucrose flavor spokes are aligned with rectangular white cue card and chocolate flavor spokes are aligned with a triangle cue card. Reward flavors act as local cues because the animal experiences having rewards with distinct flavors at different spokes.

Each flavor is exclusively available at a specific spoke only, i.e., one spoke will receive only one particular flavor of reward. The distal cues are visual, whereas local cues, here reward flavors, have distinct gustatory and olfactory features. Moreover, there can be an inherent value-based bias within reward flavors, an incentive salience that might vary across animals.

A circular LED light mounted over the ceiling delivers the required illumination for the experiment room. A white noise generator placed centrally below the track apparatus is used to mask external noise and thus prevent any bias in behavior or/and neural activity due to any external noise. The experimental session as described below begins once the tetrodes reach the target region (here, CA1), and the animal was trained to complete 15-20 laps per session. Some tetrodes were adjusted each day to maximize the number of clusters recorded.

### Experiment 1: Reward (Flavor) - Distal Cue Double Rotation

Each day of the experiment is comprised of five sessions where three standard(STD) sessions were separated by two mismatch(MIS) sessions. The configuration between the local cues (here, distinct reward flavors) and the distal cues in the STD sessions was preserved the same way as in the training sessions. In STD sessions, the banana flavor spoke is aligned (0°) to the stripe cue card, sucrose flavor spoke is aligned (0°) to the white cue card, and the chocolate flavor spoke is aligned (0°) to the triangle cue card. In the first mismatch (MIS1) session, the track was rotated 60° CCW, and the distal cues were rotated 60° CW. This change in association lead to a local-distal cue conflict where the configuration between the local cue (here, reward flavors) and the distal cues are changed. In MIS1, banana flavor spoke is aligned (0°) to triangle cue card, sucrose flavor spoke is aligned (0°) to stripe cue card, and chocolate flavor spoke is aligned (0°) to white cue card. All STD sessions have the same configurations. In the second mismatch session (MIS2), the track was rotated 120° CCW, and the distal cues were rotated 120° CW. This rearrangement leads to a distinct configuration where banana flavor spoke is aligned (0°) to white cue card, sucrose flavor spoke is aligned (0°) to triangle cue card, and the chocolate flavor spoke is aligned (0°) to stripe cue card. Animals ran 15 laps in each session. There is no waiting time for the animal to receive the reward. Five animals were used in this experiment.

### Experiment 2: Reward(flavour)+texture – distal cue double rotation

The experimental paradigm is similar to that of experiment 1, except that the track is textured. The circular track has three different types of textured surface, each covering 1/3 of the track are as follows, striped surface (black and white), black surface, and the rubber surface (blue colored). The center of the striped surface of the track is aligned with the banana flavor spoke, the center of the black surface is aligned with the sucrose flavor spoke, and the center of the rubber surface is aligned with the chocolate flavor spoke. The alignment of the distal cue and reward spoke is the same as in experiment 1 in STDs. The changed configuration in mismatch sessions is also the same as that of experiment 1. Only the alignment between the textures and the spoke (reward flavor) was maintained when the track was rotated. The number of sessions was also the same as that of experiment 1 where 3 STD sessions were separated by 2 MIS sessions. The STD configuration is changed to create a local-distal cue conflict by rotating the textured track and reward flavors (CCW) and the distal cues (CW) by either 60° or 120° in MIS1 and MIS2 conditions, respectively. A total of three animals were used in this experiment. Out of three animals, two animals were used after the completion of experiment 1. The other animal was only used for this experiment.

### General Experimental Procedure

At the start of the experiment, the rat was carried by the experimenter into the behavioral room inside a covered cardboard box to disrupt the animal’s sense of direction. The rat was then transferred on a pedestal placed at the center of the track. The tethers were connected to the EIB boards via the head stage. The rat was released at a random location on the track in every session. There is no waiting period for the rat as pellets were delivered before the rat reaches the spoke. After the completion of 15 laps, the rat was disconnected and transferred to the box for disorientation. During this period, distal cues and the track were rotated to change the configuration. We repeated the same procedure for both experiments.

After completing all sessions on an experimental day, the track and the spokes were wiped with 70% ethanol to remove any olfactory traces that could potentially bias the behavior and neural representation on the next day of recording. Rat droppings during experimental sessions, if any, were removed with tissue paper.

### Data Analysis and Statistical Tests

The data analysis and statistical tests were performed in Matlab (MathWorks R2019a), Circular Statistics (Oriana, Kovach Computing Services, UK), and Graph Pad Prism as described below.

### Isolation of single-units

All data analysis, including spike sorting, was performed offline. Single units from tetrode recordings were isolated using custom-written spike-sorting software (Winclust, Siegel *et al*., 2008; Sharma *et al.,* 2020). Cells were manually isolated based on the spike parameters such as peak amplitude, energy, peak-to-valley amplitude recorded on four tetrode channels. Based on the cluster’s distance from the background, waveform shape, inter-spike interval, and potential overlap between neighbouring clusters present, isolation quality was rated on a scale ranging from 1 to 5 (1- very good, 2-good, 3-fair, 4- marginal, 5-poor). All units rated ’fair’ and above are used for analysis. Marginal and poor clusters were excluded from the analysis. The cluster ratings were independent of the spatial property of the unit isolated.

### Place cell Identification

We used the following criteria to classify place cells:

1. Mean speed filtered firing rate in at least one session between 0.05 – 10 Hz.
2. Have a statistically significant(p<0.05) spatial information score >0.5 in at least one of the sessions on an experimental day.

Spatial information score indicates the amount of information carried about the rat’s spatial location by a cell (Skaggs *et al*., 1993). It was calculated according to the formula written described below.

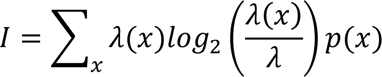

Where I is the spatial information score, *λ*(*x*) is the mean firing rate in each pixel (position), *λ* is the mean firing rate of the cell, p(x) is the probability of the rat to be in the pixel x ( occupancy time in pixel (x)/ total occupancy time.). Spatial information score was computed on a smooth speed filtered rate map (> 1cm/s).

### Rotational Correlation

To quantify the change in the firing field of place cells in the hippocampus, we conducted rotational correlation analysis on the ensemble recorded on each day of the recording. Two-dimensional occupancy and spike data were linearized to obtain a one-dimensional firing rate array of 72 bins each (5° each per bin). One-dimensional rate arrays were constructed by dividing the number of spikes fired when the rat was in a particular bin by the amount of time spent (in seconds) in that specific spatial bin on the circular track. We used a speed filter of 1 cm/sec. We have quantified the rotation of individual cells between STD sessions or between STD and MIS sessions using Pearson’s product-moment correlation between linearized firing rate arrays of a cell between two corresponding sessions. We do this procedure 71 times by shifting the session being correlated by one bin increment (5°) to obtain shifted correlation (Neunuebel *et al*., 2013, Sharma *et al.,* 2020). The angle at which maximum correlation is obtained is regarded as the amount of rotation of the place field between the sessions. Circular statistical tests were conducted both on the ensemble as well as population data (grouped) to calculate the angle and length of the mean vector using Oriana Software (Kovach Computing Services, UK). The angle of the vector represents the mean angle of rotation of the cells in the ensemble/population (grouped). The mean vector length (MVL) determines the compactness of the distribution of angles of the cells in the ensemble/population (grouped). It is inversely proportional to the variance of the distribution around the mean.

### Cluster Optimization

To identify any patterns in the mean angle of orientation across ensembles in different comparisons, we used the k-means elbow method to determine the optimum number of clusters based on the mean vector length and mean angle of rotation of all the ensembles. The optimum cluster number is found to be three across all the comparisons. We then used the k-means algorithm to group the ensemble based on their mean direction and mean vector length into three different clusters.

### Population Correlation Analysis

To measure the population activity of the hippocampus, 2D spatial correlation matrices were created from firing rate vectors (1D) at each position on the track (Neunuebel *et al*., 2013, Sharma *et al*., 2020). Population Correlation matrices for STD vs. STD and STD vs. MIS sessions were created for the grouped and pooled cells. Firing-rate arrays (1° bin) of pooled cells and grouped cells were separately combined to generate an N x 360 matrix where N is the number of cells, and 360 is the total number of positional bins. A Pearson’s product-moment correlation analysis was performed between the standard session matrix and subsequent standard or mismatch session to create a 360 x 360 correlation coefficient matrix. The population correlation matrix comprises Pearson correlation values acquired from bin-wise correlation of the 1D firing rate arrays. For STD vs. STD and STD vs. MIS comparisons, higher correlation angles would be expected at the angle equivalent to the rotation of the preferred direction of firing fields. Population responses were quantified by transforming 2D correlation matrices to 1D polar plots by calculating the average correlation of pixels in each of the diagonals in the correlation matrices. The angle that showed the maximum correlation is regarded as the amount of rotation of the population of cells between two sessions.

### Map Rotation Analysis

To identify the structure of the spatial map, firing rate arrays (1° bin) were pooled for all the cells for a particular STD session of a day. For subsequent MIS or STD sessions, 1D polar plots were created from firing rate arrays for each cell in the ensemble, and the mean direction of each cell was calculated. We then calculated the mean direction of the ensemble using the Circular Statistics Toolbox, Matlab (Berens,2009). All the firing rate arrays on that particular day were shifted according to the round-off value of the mean direction of the ensemble. For example, if the mean direction of an ensemble is 62° in STD1 vs MIS1 comparison, firing rate array of all the active cells in that particular ensemble (from MIS1) is shifted by 62°. This procedure was repeated for all the ensembles, and all shifted firing rate arrays were pooled. Shifted population correlation matrices of STD vs. shifted STD and STD vs. shifted MIS comparisons were created. The creation of population correlation matrices and quantification of the matrices using a polar plot is the same as that explained in the Population Correlation Analysis. If the hippocampus maintains a single map, we could see a single band of high correlation at the diagonal as the shifted map aligns at the diagonal in all the comparisons. To quantify the shifted population correlation matrix, the polar plot was calculated from 2D correlation matrix, and mean vector length (MVL) of the polar plot was calculated. We used the shuffling method to analyse whether the single map obtained is out of chance. In this method, we shuffled the mean direction of 33 ensembles in each comparison (shuffling ensemble mean directions of 33 days so that each ensemble is shifted by the mean direction of any random ensemble). All active neurons in a particular ensemble were shifted to the round-off value of a random shuffled mean direction. We performed this shuffling procedure for all the ensembles, pooled the neurons, and performed the map rotation analysis of shifted STD/MIS sessions based on shuffled mean directions. The shuffled population correlation matrix was thus created. To quantify the shuffled population correlation matrix, the polar plot was calculated from 2D correlation matrix, and mean vector length (MVL) of the polar plot was calculated. This procedure was repeated 1000 times, and the 99^th^ percentile of the MVL of the shuffled correlation matrix was calculated.

### Spatial cross-correlation

As one of the characteristic features of attractor dynamics in the spatial neural network is the maintenance of spatial offset between place fields of co-recorded cells, the spatial offset between co-recorded place cells were compared across STD and Mismatch sessions by modifying the analysis of Basset *et.al*, 2018, in such that, spatial cross-correlation was performed on the entire time duration in a session (Sharma *et al*., 2020), as described below. Two-dimensional spike and occupancy data were linearized to produce 1D firing rate arrays of bin size 5° (total 72 bins). The firing rate arrays were divided by their peak value to create normalized firing rate arrays. The Spatial Cross-Correlation (SXC) arrays were created by calculating the Pearson’s Product moment correlation of normalized firing rate arrays between co-recorded cell pairs in a session and its shifted correlation (5° increments) which was repeated 71 times. The spatial offset angle that gave maximum SXC was regarded as the peak correlation angle values. By pooling all cell pairs across sessions and rank order based on the peak correlation angle in STD1, SXC matrices were created for all sessions. The rank order was maintained in all other sessions. To quantify the spatial offset of SXC matrices, polar plots for each SXC cell pair (72 bins) were created, and the mean direction of the polar plot was determined using a MATLAB-based circular statistics toolbox (Berens, 2009). The mean direction value of a cell pair in a particular session represents its spatial offset in that session. The coupling between cell pairs in the population was quantified by correlating the spatial offset values (after correcting for -180°/180° transition values) between the STD vs. MIS and STD vs. STD sessions by calculating r-squared values and their statistical significance.

### Histological Analysis

Marker lesions were made on a selected few tetrodes by passing current (10 μA for 10 seconds). Rats were perfused transcardially on the following day using 4% formaldehyde solution. After perfusion, the brain was extracted and preserved in 30% sucrose-formalin solution till it sank.

Later, brains were sectioned (Coronal plane, 40 μm sections) and mounted left to dry. Sections were then stained through the Nissl stain procedure using 0.1% Cresyl Violet. Images of the sections were acquired using Leica DFC265 digital camera connected to Leica M165-C stereo microscope. The tetrodes on the sections were identified using tetrode arrangements in the Microdrive. Depth reconstruction was conducted on each tetrode track to identify the brain region from which electrophysiological recordings were conducted each day, based on the tetrode tip distance and considering the shrinkage factor (assuming 15% shrinkage occurred to histological processing)

## Results

We studied the dynamics of the spatial representation in the hippocampal CA1 population when the animal experienced a change in the association between reward flavor (local cues) and distal cues in the environment. We have recorded a total of 680 neurons from six rats in this study, while the rats ran CW on a plain track (591 neurons, experiment 1, 5 rats) and a textured track (89 neurons, experiment 2, 3 rats) both in the presence of distal cues. **Fig. 1A** shows the representative examples of tetrode localization in the nissl stained coronal sections of the hippocampal CA1.

**Figure 1:**
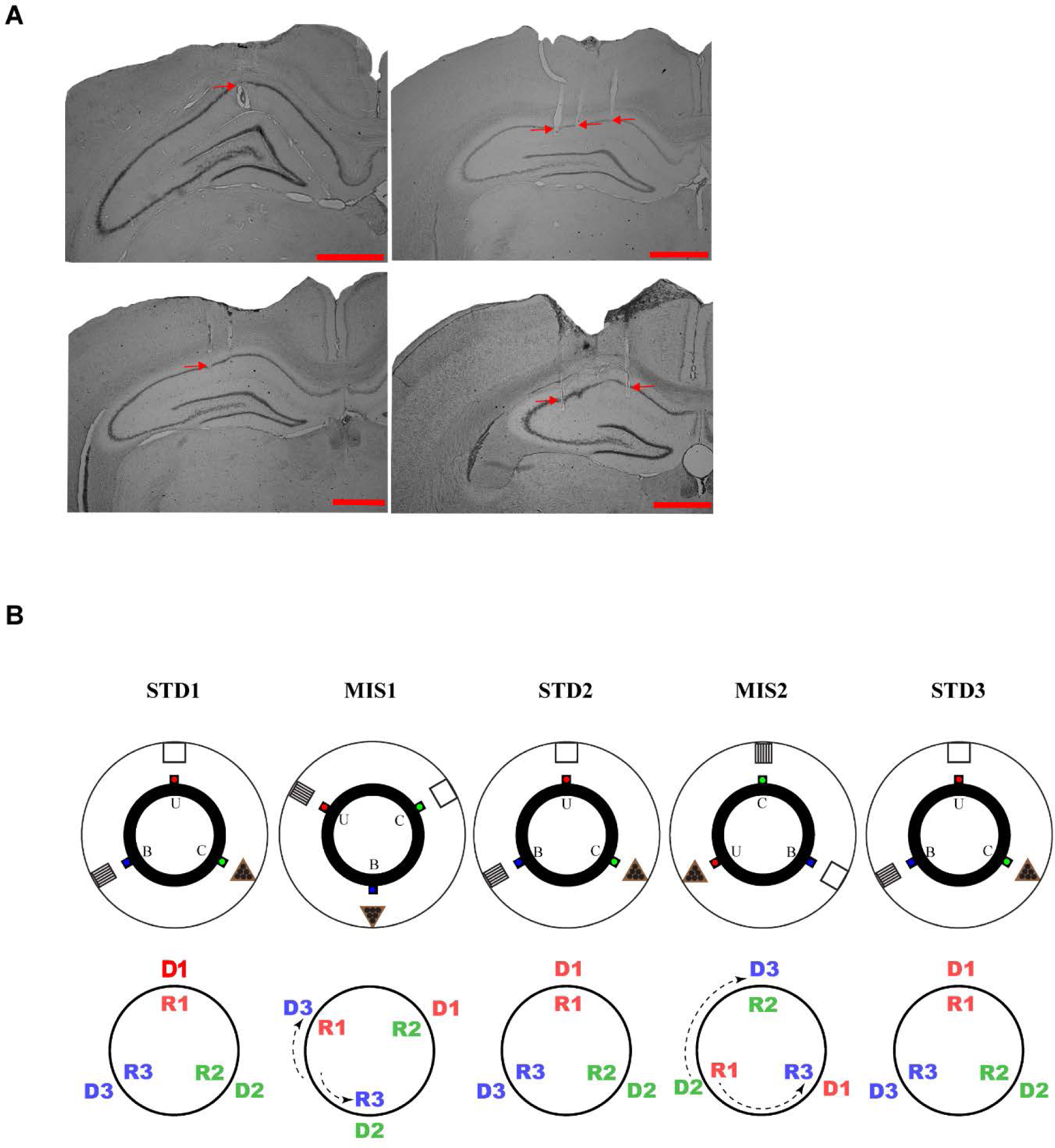
Schematic representation of the reward flavor-distal cue double rotation paradigm. **A.** Representative examples of Nissl stained coronal sections of the rat brain showing tetrode tracks (indicated by arrows) in the CA1 region of the hippocampus. (Scale bar=1 mm) **B.** Top row shows the schematic representation of the reward flavor-distal cue double rotation paradigm. Blue dot indicates the spoke where banana flavor pellets(B) were provided, red dot indicates the spoke where Sucrose pellets(U) were provided and green dot indicates the spoke where Chocolate flavored pellets (C) were provided. Bottom row shows the graphical representation of the rotation of reward flavors and distal cues. R1, R2 and R3 indicates spokes that provided sucrose, chocolate flavored and banana flavored pellets respectively. D1, D2 and D3 indicates the respective distal cues aligned with R1, R2 and R3. D1 and R1 is shown in red, D2 and R2 is shown in green, and D3 and R3 is shown in blue. Arrow inside and outside the circle indicates the direction of rotation of the track (reward flavors) and distal cues.

### Experiment 1: Reward (Flavor) - Distal Cue mismatch experiment

We used a novel, modified version of the double rotation paradigm (Knierim, 2002) to study the dynamics of the spatial representation in the hippocampus when the association between distal cues and reward flavors is dynamically changed. We recorded 591 CA1 neurons from 5 rats in this experiment (obtained over 33 recording days, five sessions each).

In this experiment, rats were trained to run CW on a plain black track with three spokes (black) separated 120° apart from each other. Three different flavors of pellets, namely Sucrose, Chocolate, and Banana flavoured pellets (Bioserv Dustless Precision pellets, 45 mg), served as rewards and were manually delivered at each specific spokes. In this experiment, reward flavors were considered as local cues as the animal directly experiences these cues. Each spoke (with distinct reward flavors) was associated with visually distinctive distal cues (made of cardboard) hung on the surrounding black curtain at the periphery of the experimental room. The experimental paradigm is shown in **Fig. 1B** and its graphical illustration in **Fig. 1C**. Each day of the experimental session comprises three standard sessions(STD) separated by two mismatch sessions(MIS). In the STD session, the white cue card was aligned with the sucrose spoke, the triangle cue card was aligned with the chocolate spoke, and the striped cued card was aligned with the banana spoke (see Fig. 1B). In the MIS session, the track was rotated CCW, and distal cues were rotated CW by either 60° (MIS1) or 120° (MIS2), leading to a change in the association between reward flavors and distal cues.

Previous studies had shown that hippocampal ensemble showed heterogeneous representation when local textures on the track and distal cues were rotated. In these studies, simultaneously recorded place cells rotated their place fields CW with distal cues, CCW with local cues (textures), and some remapped in a particular session (Knierim, 2002; Lee *et a*l., 2004; Yoganarasimha *et al*., 2006). On the other hand, when reward flavors and distal cues were rotated in this experiment, we observed a unified response where all the cells dynamically oriented to only one cue type simultaneously in a particular session. **Fig. 2** shows distinct dynamic orientations of representative example place fields to distal cues, reward flavors, or following arbitrary rotation (not following either distal cues or reward flavors) from three different ensembles (five example cells each) from three different rats. In ensemble-1 (**Fig. 2A**), all the cells maintained their place fields across STD sessions on the track. These results suggest a stable hippocampal representation across the STDs. During mismatch sessions, the representation was oriented to the reward flavors (local cues) where CA1 place fields rotated CCW in the mismatch sessions, following the rotation of reward flavor cues on the track. All the cells in this ensemble (ensemble-1) showed the same responses across sessions. (for the rate map of all the cells in ensemble-1, please refer to **Supplementary Fig. 1-1**). In ensemble-2 (**Fig. 2B**), all the cells maintained their place fields between STD1 and STD2 (only five cells were recorded in this experimental day from two tetrodes). However, in STD3 it showed a different orientation by rotating with reward flavors. In MIS1, all the place fields rotated CW following the rotation of distal cues. However, in MIS2, the place fields were not controlled by either distal or local cues. Instead, all the place fields maintained their position as same in STD2. These results show an overall distinct representation of ensemble-2 compared with ensemble-1 where MIS sessions follow distinct orientation and all the STDs do not maintain stable place fields. But all the cells showed a unified response by all cells following only particular cues across sessions. In ensemble-3 (**Fig. 2C**), STD1 and STD2 did not show a stable representation, whereas STD2 and STD3 showed a stable representation. This result indicates a dynamic change in orientation across STDS as place cells did not fire in the same location in all the three STDs. In MIS1, place fields followed neither distal nor reward flavor. Instead, it flips its firing field 180° to an arbitrary angle (in conjunction with the position with respect to spokes/cues). In MIS2, all the place fields rotated coherently following the rotation of reward flavors. This response is different from that observed in ensemble1 and ensemble2 (for the rate maps of all cells in the ensemble, refer to **Supplementary Fig. 1-2**). Results from **Fig. 2** suggest that hippocampal representations are dynamic in a reward-distal cue mismatch experiment. In some sessions, it was oriented with local cues (reward flavors), and in some sessions, it may orient with distal cues or may exhibit an arbitrary orientation. But even under such dynamic conditions, the hippocampus showed a unified response where all the simultaneously recorded CA1 neurons rotated their place fields towards one type of preferred cue in a particular session (either distal cues or reward flavors but never both simultaneously). We observed such a dynamic change in cue orientation across most sessions and experimental days.

**Figure 2:**
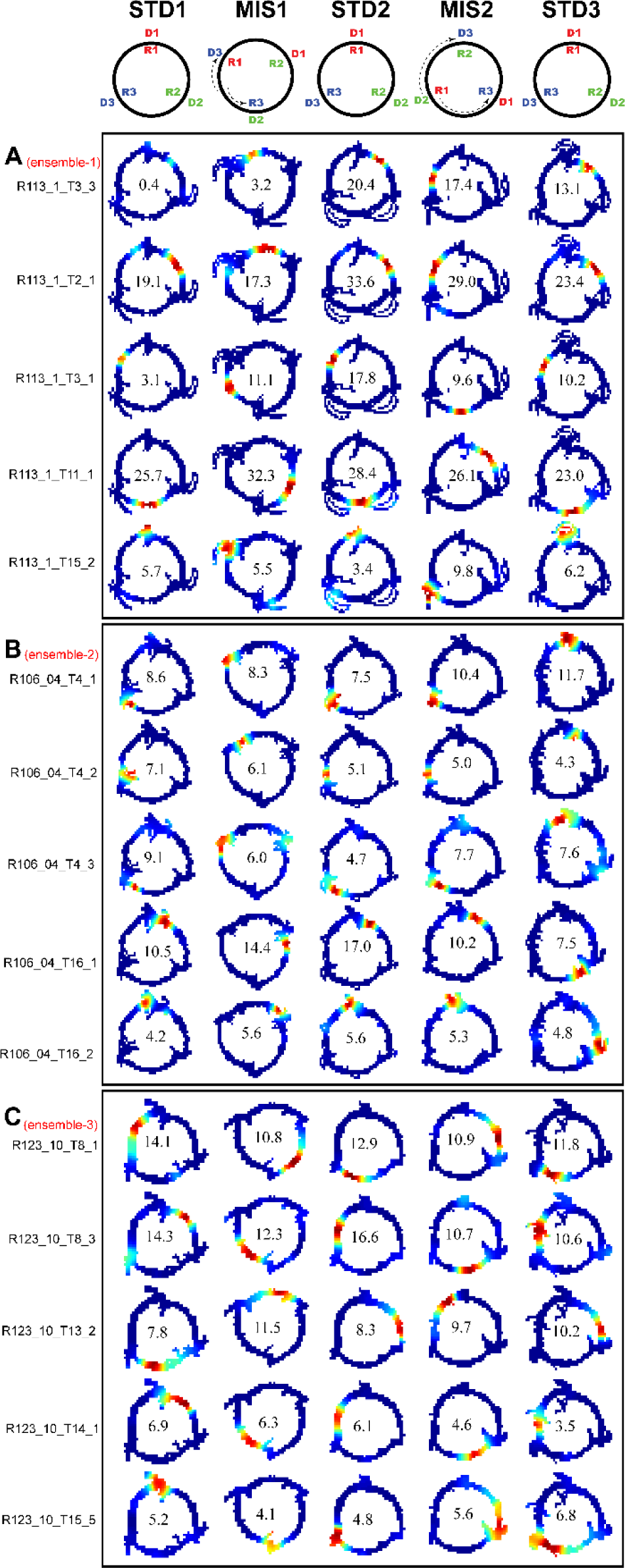
Representative examples of subset of neurons from different ensembles. **A-C** shows rate maps of five neurons each from three different ensembles (from different rats). Left side of the plot shows the IDs of each neurons. **A.** Ensemble-1: Representative examples of simultaneously recorded CA1 firing rate maps in an ensemble across standard and mismatch sessions. In mismatch sessions all the place fields followed the rotation of track based reward flavor CCW. (For the rate maps of all the cell in this ensemble refer to **Supplementary Fig 1-1**.) Ensemble-2: Representative examples of simultaneously recorded CA1 firing rate maps in another ensemble across standard and mismatch sessions. In the first mismatch session(MIS1), all the place fields followed the rotation of distal cues. In the second mismatch session(MIS2), place fields did not follow either cues. In the subsequent standard session, all the place fields follow the rotation of reward flavors. **C.** Ensemble-3: Representative examples of simultaneously recorded CA1 firing rate maps in another ensemble across standard and mismatch sessions (For the rate maps of all the cell in this ensemble refer to **Supplementary Fig 1-2**.). In the first mismatch session(MIS1), neither reward flavors nor distal cues controlled the place fields. In the second mismatch session(MIS2), all the place fields were controlled by reward flavors. The number inside the firing rate maps indicates the peak firing rate (in Hertz (Hz)). The rate maps are color coded, with red color denoted by >90% of the peak firing rate, no firing is indicated by blue and successive gradients in the firing rates are shown with intervening colors in the spectrum.

To quantify the rotation of place fields in the ensembles between STD vs. MIS and STD vs. STD comparisons, rotational correlation analysis was performed on all the cells that showed activity in the respective comparisons in each ensemble. In this procedure, we linearized two-dimensional circular track data to produce one dimensional firing rate array (5° bin, 72 bins) for each cell for all sessions. The angle of peak correlation was calculated for all simultaneously recorded cells in both STD vs. MIS comparisons and STD vs. STD comparisons. Circular Statistical analysis was performed on each ensemble across all the comparisons to study the distribution of peak correlation of all the angles in the ensembles. In this analysis, the angle of the mean vector and the mean vector length (MVL) of the distribution of all the comparisons were calculated for all the ensembles. The angle of the mean vector for the ensemble constitutes the circular mean of the angle of rotation of the ensemble. The MVL of the ensemble is inversely proportional to the variance of the distribution of individual cell rotation angle in that ensemble (based on its peak correlation). Thus, the MVL values indicate the compactness of the distribution of rotation angles of individual cells in the ensemble, which is considered a measure of coherency (Leet *et al.,* 2004; Yoganarasimha *et al.,* 2006).

**Fig. 3A** shows the box plot of the MVL across STD vs. MIS and STD vs. STD comparisons for all the ensembles. The average MVL across ensembles in the pooled STD1 vs. MIS1, STD2 vs. MIS2, STD1 vs. STD2, and STD2 vs. STD3 comparisons are 0.921±0.015, 0.933±0.014, 0.907±0.015, and 0.930±0.016 respectively (mean ± S.E.M.). This suggests that the peak angles of correlation of individual neurons in each ensemble across all the comparisons were clustered in one particular direction (in a particular session). In all the comparisons across all the ensembles, the mean vector length is high and statistically significant ((> 0.8, Lee *et al*., 2004; Rayleigh test: p<0.01, in all the comparisons across all the ensembles), indicating a coherent structure in the orientation. **Fig. 3B** shows the distribution of the mean direction of ensembles of individual rats (color-coded) across STD and MIS comparisons. This result shows that all the rats exhibited dynamic orientation to cues as observed in **Fig. 2**. In STD vs. STD conditions, ensembles show distinct orientations (24% CCW (∼ -120°), 24% CW (∼ 120°), 52% Stay (∼ 0◌_ۣ_ °) in STD1 vs. STD2 comparison; 30% CCW (∼ -120°), 27% CW (∼ 120°), and 43% Stay (∼ 0◌_ۣ_ °) in STD2 vs. STD3 comparison). Stay indicates the ensemble mean direction data points that showed stable place fields across STD sessions. These results indicate that in STD comparisons, representations are overall unstable indicating a dynamic change in the orientation in subsequent STD sessions.

**Figure 3:**
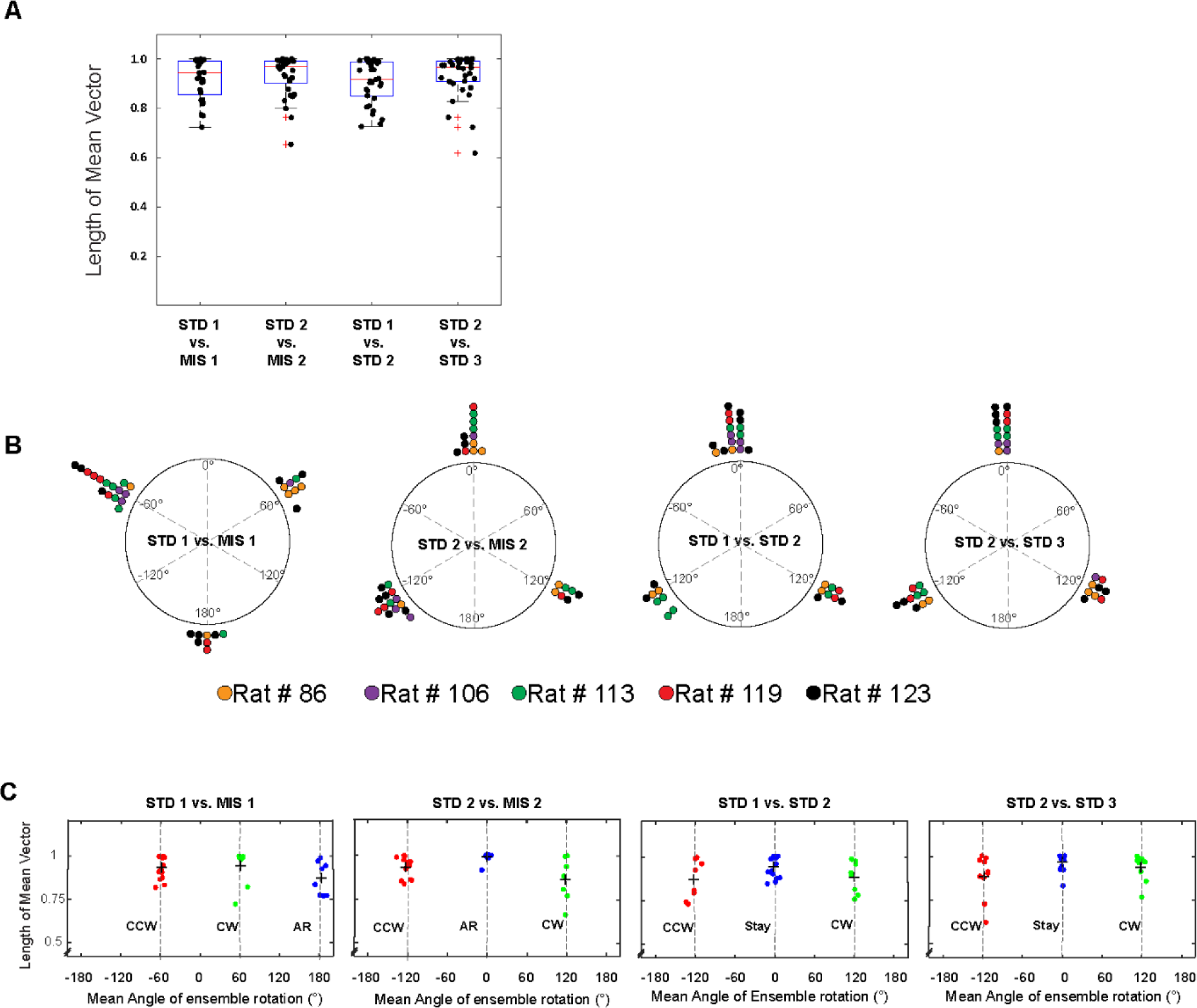
Mean Vector Length Distribution and dynamics of place cell orientation. **A.** Box plot showing the distribution of mean vector length of all the ensembles across standard and mismatch sessions. + sign indicates outliers. Each dot represents the mean vector length of an ensemble. **B.** Distribution of mean direction of the rotation of all the ensembles across standard and mismatch sessions of individual rats (color-coded). Each dot represents the angle of mean vector (mean direction) of an ensemble. **C.** Scatter plot of the mean direction and mean vector length of all the ensembles across standard and mismatch sessions. Red, Green and Blue dots show the clustering of the data points based on k-means elbow method. ‘+’ indicates the centroid of each clusters. In standard vs mismatch comparisons, red cluster signifies the ensembles that followed the rotation of CCW reward flavors (CCW), green cluster signifies the ensembles that followed the rotation of CW distal cues(CW) and blue cluster signifies the ensembles that rotated at an arbitrary angle(AR), not following either reward flavors or distal cues. In standard vs standard comparisons, red cluster signifies the ensembles that followed the net CCW rotation (CCW), green cluster signifies the ensembles that followed the net CW rotation (CW) and blue cluster signifies the ensembles that showed a stable representation between standard sessions(Stay). Grey dotted line denotes the angle of separation of spokes/cues (120°).

However, many ensembles showed more stable response(Stay) in STD vs. STD comparisons compared to CCW and CW alignment. Overall, In STD vs. MIS conditions, more ensembles oriented toward the reward flavors than the distal cues (48% CCW (∼ -60°), 27% CW (∼ 60°) and 24% arbitrary rotation (∼ 180°) in STD1 vs. MIS1 comparison, 42% CCW (∼ -120°), 24% CW (∼ 120°) and 34% arbitrary rotation (∼ 0°) in STD2 vs. MIS2 comparison). This result shows that the orientation of ensembles in a cue mismatch environment is essentially dynamic and slightly biased towards the reward cues for the orientation. Out of 33 days of recording, in six days, the rotation was not dynamic and all the neurons in that ensembles followed one particular cue only. On other days (across rats), the ensembles showed a dynamic shift in orientation (in at least one session) from one cue to another or shifted to an arbitrary angle. Even during some standard sessions, we observed a dynamic shift in the orientation leading to unstable place fields. However, even during this instability, the place fields coherently rotated away from the initial location.

As individual ensembles across comparisons showed a coherent structure with a high mean vector length (from **Fig. 3A**) and also showed distinct orientation across sessions (from **Fig. 3B**), we investigated if any underlying pattern exists in these ensembles. We used k-means algorithm and implemented the elbow method to optimize the number of clustered ensembles obtained based on the mean vector length and mean direction of each ensemble across sessions. Through this method, the population response of distinct clusters of ensembles (based on coherency and orientation) can be evaluated. In all the comparisons, the desirable number of clusters was found to be three. We then used k- means clustering to obtain three different clusters of ensembles across comparisons. **Fig. 3C** shows the scatter plot of the ensemble mean vector length and mean direction with three clusters across STD vs. STD and STD vs. MIS comparisons. Of the three cluster groups aligned in STD vs. MIS comparisons, the cluster which is aligned to the CW rotation of distal cues was named CW group, the cluster which is aligned to the CCW rotation of reward flavors was named CCW group and the cluster oriented to an arbitrary angle (not following CW and CCW rotation) were named as Arbitrary Rotation (AR) group. Thus, each group has ensembles orientated towards a specific orientation where CCW and CW groups were aligned to CCW and CW rotation of reward flavor and distal cues, respectively. The AR group is located almost equidistant from CCW and CW group (∼120°), indicating the spacing of the orientation of the ensembles based on the angle of separation between the spokes. In STD vs. STD comparison, CCW cluster and CW cluster were aligned to the net CCW and CW rotation between STD sessions indicating the dynamical shift in cue preference between STD sessions. The other group was named Stay group because the representation between the STD sessions was stable in all the ensembles in that group by staying in the exact location aligned with the same distal cue – reward flavor association and showing the ensembles’ clustering near 0°.

The cluster groups obtained organized the ensembles into three groups in all STD vs. STD and STD vs. MIS comparisons, and rotational analysis was performed in the entire neural population in each group. All the active neurons were pooled together from each ensemble in a particular cluster group from their respective STD vs. STD and STD vs. MIS comparisons. Circular statistics were performed on all ensemble data from each group and the distribution was plotted as shown in **Fig. 4**. The polar plots depict the data distribution from all the rats in each group for STD vs. MIS comparisons (**Fig. 4A and 4B**) and STD vs. STD comparisons (**Fig 4C and 4D**). The circular statistics of STD vs. MIS comparisons in the CW group show that the angle of the mean vector moved to the amount of rotation of the distal cues in MIS sessions. Also, the angles show clustering with a significantly high mean vector length (> 0.8, Lee *et al*., 2004) indicating a coherent rotation (STD1 vs. MIS1(CW) =0.912, STD2 vs. MIS2 (CW)=0.87, Rayleigh test, p<0.0001 for both the comparisons). In STD vs. MIS comparisons in the CCW group, the angle of the mean vector deviated to equal to the amount of rotation of the reward flavors. Here also, the angles show a significant clustering with high mean vector length in both the comparisons indicating a coherent rotation (STD1 vs. MIS1(CCW) =0.918, STD2 vs. MIS2(CCW)=0.914, Rayleigh test, p<0.0001 in both the comparisons). Further, STD vs. MIS comparisons in the AR group also show that the angle of the mean vector deviated equally from CW and CCW rotation. Even in this group, the angles show a significant clustering with a high mean vector length (STD1 vs. MIS1(AR) =0.874, STD2 vs. MIS2(AR)=0.980, Rayleigh test, p<0.0001 in both the comparisons).

**Figure 4:**
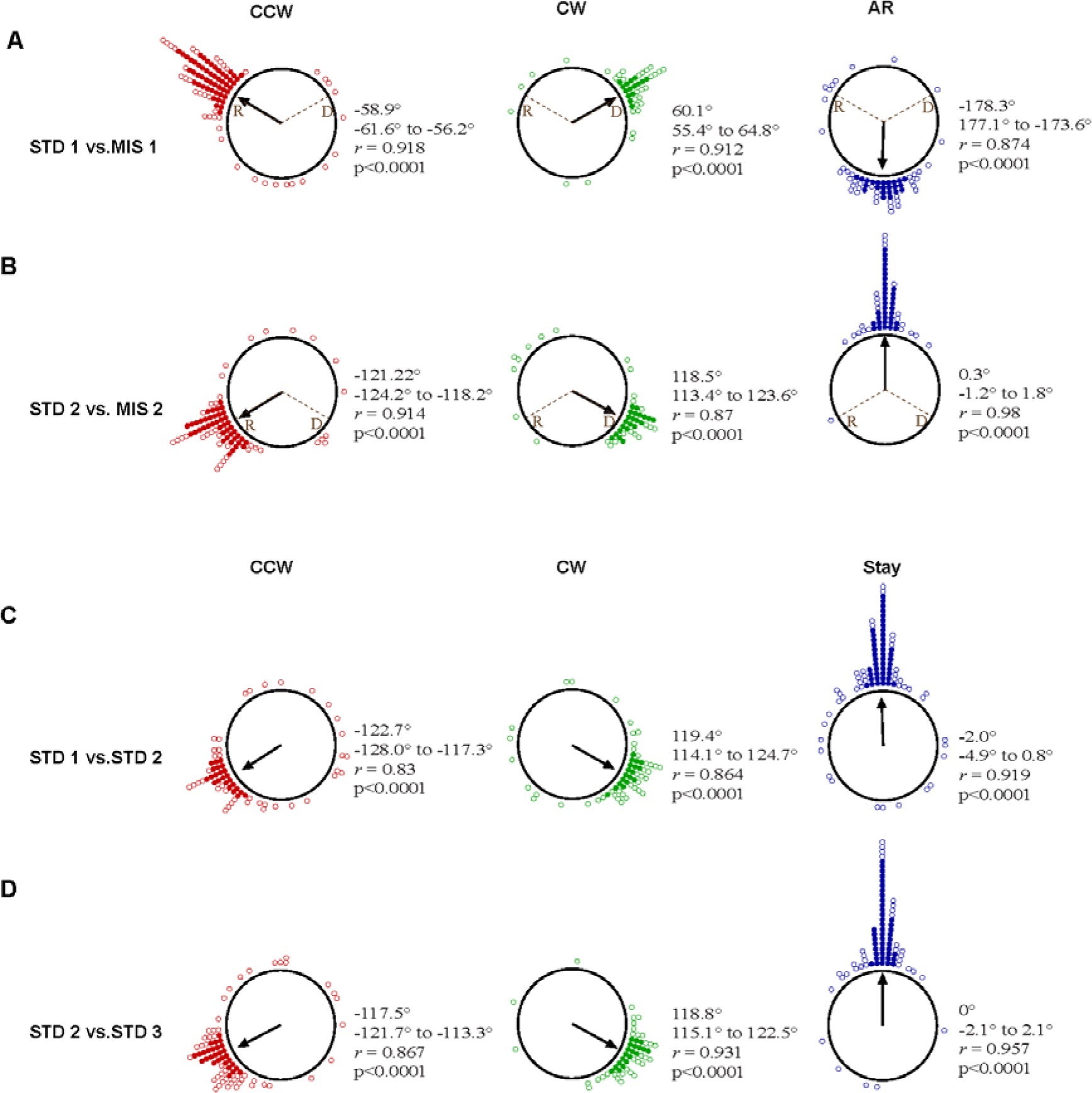
Dynamic and coherent representation in the CA1. **A-B:** The amount of rotation of firing fields of all the active place cells in all the ensembles in the CCW group (red circle), CW group (green circle) and AR group (blue circle) between STD vs. MIS sessions, represented around the circle. **C-D:** The amount of rotation of firing fields of all the active place cells in all the ensembles in the CCW group (red circle), CW group (green circle) and Stay group (blue circle) between STD vs. STD sessions, represented around the circle. The direction of arrow represents the mean angle of rotation of population of neurons in each group and length of the arrow signifies the compactness of the distribution around the mean angle. Brown line indicate the rotation angle of reward flavors (R) and distal cues (D) in MIS sessions. (circle; open=1 cell, filled=5 cells). Values of mean angle of the distribution, 95% confidence interval, length of the mean vector (r) and its significance level (p) are shown near to each plot.

In STD vs STD comparisons in the Stay group, the place fields were significantly clustered near 0° and have significant mean vector length (STD1 vs. STD2 (Stay) =0.919, STD2 vs. STD3 (Stay) = 0.957, p<0.0001 in both the comparisons). In other groups, the cells were clustered around in either CW (STD1 vs. STD2 =0.864, STD2 vs. STD3=0.931, Rayleigh test, p<0.0001 in both the comparisons) or CCW (STD1 vs. STD2 =0.83, STD2 vs. STD3=0.867, Rayleigh test, p<0.0001 in both the comparisons) direction. These results suggest that across all comparisons between STD vs. STD and STD vs. MIS comparison in different groups, place fields rotated coherently, indicating a dynamic and coherent population activity.

### Population coherence in the hippocampus

In the **Fig. 4**, rotational correlation analysis was performed at the level of its correlation peak, and it shows a coherency in the rotation of place fields in all the grouped ensembles. This analysis alone would not yield an in-depth understanding of location-specific changes in the hippocampal population in STD vs. STD and STD vs. MIS comparisons. Therefore, we conducted population correlation analysis to measure the extent of coherency across each positions on the track. Here, population responses between STD vs. STD and STD vs. MIS comparisons were measured through 1° bin-wise correlation of linearized firing rate arrays of all the neurons at each position on the track (**Fig. 5**) (Neunuebel et al., 2013; Sharma et al., 2020).

**Figure 5:**
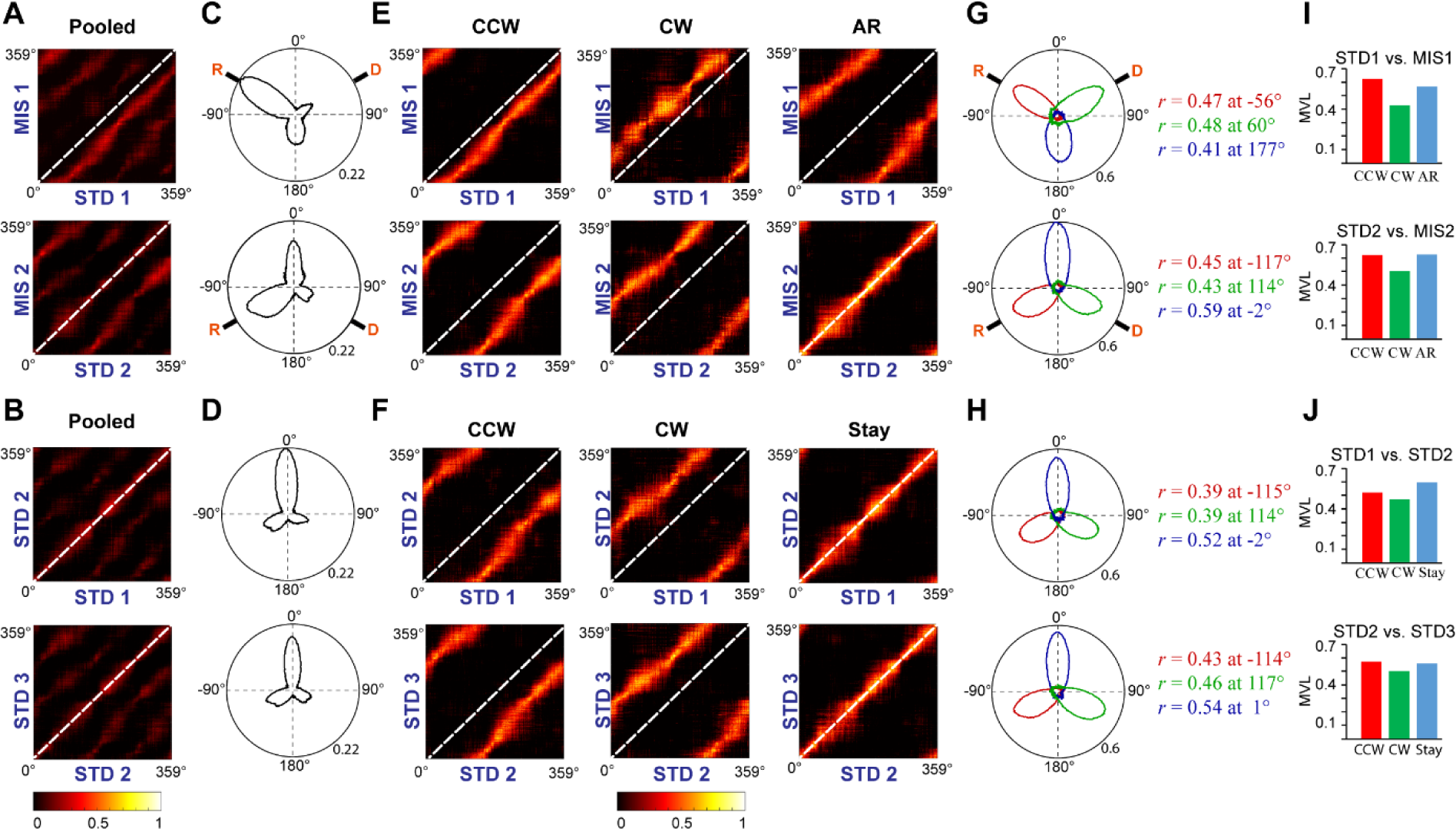
Population response in the hippocampus. **A –B**: Correlation matrices from population firing rate vectors of all the cells pooled from each ensemble at each position on the track between STD vs MIS(A) and STD vs STD sessions(B). **C-D:** The hippocampal population activity from A and B are represented as polar plots in **C** and **D** respectively. **E-F**: Grouped correlation matrices from population firing rate vectors of all the cells present in each cluster of ensembles at each location on the track between STD vs MIS (CCW, CW and AR) and STD vs STD sessions (CCW, CW and Stay). **G-H:** The hippocampal population activity of different groups between STD vs MIS sessions (CCW (red), CW(green), AR (blue)) and STD vs STD sessions (CCW(red), CW(green) and Stay(blue)) are represented as polar plots (constructed from correlation matrices). The values next to the polar plot indicates the angle of peak correlation (r) of the hippocampal population at corresponding groups. **I-J:** Length of the mean vector in each groups in STD vs. MIS and STD vs. STD sessions for each of the corresponding polar plots is shown **G** and **H**.

STD vs. STD correlation matrices has multiple bands indicating that the population response of the hippocampus was unstable across different STD sessions. STD vs. MIS correlation matrices also has multiple bands suggesting a dynamic orientation of cues across experimental days (**Fig. 5A & 5B**). To quantify the population response across groups and categories, 2D correlation matrices were transformed to 1D polar plots by calculating the mean correlation of pixels in each diagonal. The polar plot also shows the presence of three different peaks indicating dynamic orientation to three directions (**Fig. 5C & 5D**). This indicates that the population response also show dynamic orientation and thus, the population coherence could not be studied from all the neurons pooled together.

So, we used the clusters obtained from the k-means analysis and used the grouped ensembles in Fig 3C. We pooled all the active neurons from each ensemble in the respective group (as same as in Fig. 4) and bin-wise correlation was performed for all the comparisons in respective groups.

We calculated the population correlation of different orientation population responses of CCW, CW and AR groups in STD vs. MIS and CCW, CW and Stay groups in STD vs. STD comparisons. All correlation matrices within grouped STD vs. MIS comparisons have a band of high correlation in CCW, CW, and AR groups (**Fig. 5E**). In the CCW and CW groups of STD vs. MIS comparisons, the band of high correlation follows the rotation of reward flavors and distal cues, respectively. In the AR group, the band of high correlation is seen equidistant from CCW and CW bands indicating a population coherence away from CCW and CW rotation. A similar response is seen in STD vs. STD sessions, where a single band of high correlation is present in CCW, CW, and the Stay groups (**Fig. 5F**). A band of high correlation observed on the central diagonal in the Stay group indicates no change in the orientation of firing fields. In CCW and CW groups, the band shifted to CCW and CW direction indicating a shift in orientation between two STD sessions. These suggest that even though the population response of the hippocampus was unstable across some STD sessions and dynamically orient in MIS sessions, all the cells rotated as an ensemble under all the conditions, thus exhibiting a location-specific coherent population activity.

The polar plots also show the presence of single peak in all the groups across comparisons. In STD vs. MIS comparisons, the angle of peak correlation in CCW, CW, and AR group was equivalent to the rotation of track (in CCW group, -56° in STD1 vs. MIS1 comparison, -117° in STD2 vs. MIS2 comparison), rotation of distal cue (in the CW group, 60° in STD1 vs. MIS1 comparison, 114° in STD2 vs. MIS2 comparison), and arbitrary rotation angle (In the AR, 177° in STD1 vs. MIS1 session, -2° in in STD2 vs. MIS2 session) respectively (**Fig. 5G**). In STD vs. STD comparison, in the Stay group, the angle of peak correlation was near to 0° (-2° in STD1 vs. STD2 comparison, 1° in STD2 vs. STD3 comparison), indicating the stability of the place fields. In the CCW and CW group, the angle of peak correlation shifted to CCW (-115° in STD1 vs.

STD2 comparison, -114° in STD2vsSTD3 comparison) and CW directions (114° in STD1 vs. STD2 comparison, 117° in STD2 vs. STD3 comparison) (**Fig. 5H**). **Fig. 5I & 5J** shows the mean vector length of the polar plots at each group and STD vs. MIS and STD vs. STD comparisons, respectively. Also, there is no significant difference in the mean vector length across different comparisons (Kruskal-Wallis test, p=0.826), indicating similar responses in both STD vs. STD and STD vs. MIS comparisons. These results show that the hippocampus maintains a location-specific coherency despite the dynamic change in the orientations.

### Single Map in the Hippocampus

The population correlation analysis suggests that despite being unstable in some sessions in STDs, the hippocampal population maintains a coherent network. The same was seen in the STD vs. MIS comparison, where the hippocampus encodes a coherent representation of the space, even when the preference of the cues and thus the orientation of the cells are dynamically changing.

These results bring out a very fundamental question: How many spatial maps do the hippocampus employ to code for a dynamically changing reward flavour- distal cue association? To answer this question, we performed map rotation analysis using shifted firing rate arrays (modified from Kinsky *et al*., 2018). Here, shifted mismatch / STD firing rate arrays were created by shifting the linearized firing rate arrays (1°) of all the active cells in a particular day by their ensemble mean direction. All shifted sessions were pooled together to create a shifted MIS/STD session **Fig. 6(A-D**). If the hippocampus maintains a single map, we could see a single band of high correlation at the diagonal as shifted map aligns at the diagonal in all the comparisons. On the other hand, if the hippocampus exhibits more than single map, multiple bands or lack of any specific bands would be observed for the same. **Fig. 6A & 6B** shows a single band aligned at the diagonal in STD1 vs. shifted MIS1 and STD2 vs. shifted MIS2 comparison. The mean vector length is higher than the 99^th^ percentile of the shuffled distribution (MVL in STD1vs. shifted MIS1 =0.677, MVL in STD2 vs. shifted MIS2 =0.73, p<0.01). **Fig.6C & 6D** also shows a single band aligned at the diagonal in STD1 vs. shifted STD2 and STD2 vs. shifted STD3 comparison. The mean vector length is higher than the 99^th^ percentile of the shuffled distribution (MVL in STD1 vs. shifted STD2 =0.69, MVL in STD2 vs. shifted STD3 =0.73, p<0.01). All comparisons show a single band along the diagonal. To quantify the population correlation matrices, 2D correlation matrices obtained were transformed to 1D polar plots by calculating the mean correlation of pixels in each diagonal. Across all the comparisons, mean direction and the angle of peak correlation was near to 0° (3° in STD1 vs. shifted MIS1, - 1° in STD2 vs shifted MIS2, 2° in STD1 vs. shifted STD2 and 1° in STD2 vs. shifted STD3 comparisons) with the mean vector length more than the 99^th^ percentile of the shuffled distribution mean vector length (see methods). This striking result strongly suggests the use of a single coherent spatial map by the hippocampus to represent dynamically changing environmental association. This single map may be oriented to different cues in different sessions. Still, within a session, the single map orients to one particular direction thus be able to produce a single representation. This shows that despite being unstable in some STD sessions and dynamically orient to different reference frames in MIS sessions, hippocampus shows a single map by coherent rotation of its place fields assemblies.

**Figure 6:**
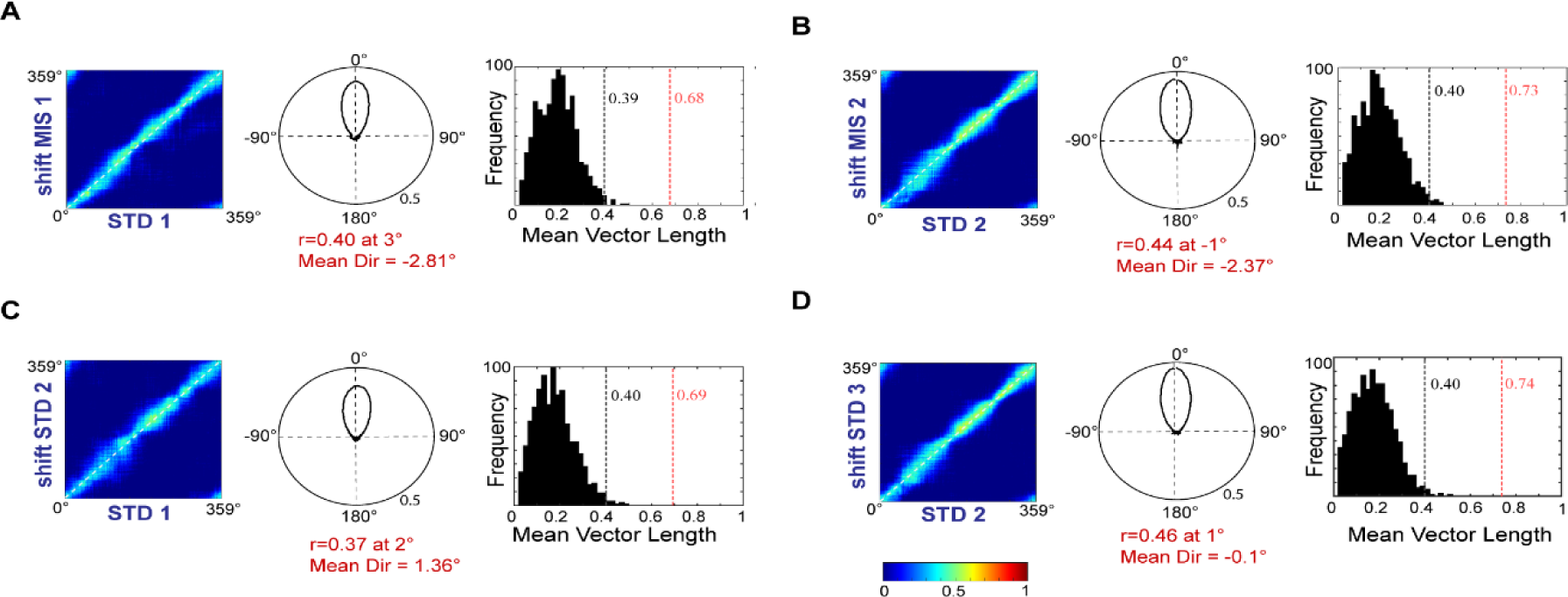
Single Spatial Map in the hippocampus. Correlation matrices created from population firing rate vectors of all the cells of STD with shifted population firing rate vectors of MIS (A, B) and STD (C, D) sessions shifted based on the mean direction of rotation of the ensemble to which each cell belong to. The polar plot next to the correlation matrices indicates the quantification of the 2D correlation matrices. The values at the bottom of the polar plot denotes the angle of the peak correlation(r) and mean direction of the polar plot (Mean Dir). The frequency histogram near to the polar plot denotes the distribution of the mean direction of the polar plots created from shifted population firing rate vector based on shuffled mean direction values of ensembles across days. The red dotted line indicates the observed mean vector length and the black dotted line indicates the 99^th^ percentile of shuffled mean vector length.

This is strikingly different from double rotation experiments where the textured track was used as local cues. In those experiments, the hippocampus is shown to have a heterogeneous representation. The observed coherency and the existence of single map in a reward flavour – distal cue double rotation paradigm pose another interesting question – How would the hippocampus representation changes when the textured tracks where added on the same paradigm. In such a condition, both reward flavors and textured track may act as local cues. Based on CA1 response on double rotation experiments with the textured track, the hippocampus may establish heterogeneous representation (Knierim, 2002; Lee *et al*., 2004, Yoganarasimha *et al*., 2006). Alternatively, presence of reward flavors may still organize the hippocampal representation as a coherent map due to the salience and value aspect of the local cues. To test these hypotheses, we carried out a different experiment (Experiment 2) to study the nature of hippocampal representation upon adding textures on the track to see if the hippocampus still maintains a coherent map as shown in experiment 1.

### Experiment 2: Reward(flavour)+texture – distal cue double rotation

In this experiment, we recorded 89 place units from the hippocampal region from 3 rats (10 days, 50 sessions). Two out of three rats were used after the completion of experiment 1 and one rat is used only for experiment 2. Here, the experimental paradigm is similar to experiment 1, except that track is not plain but has distinct surface textures (**Fig. 7A**). Similar to experiment 1, in the MIS sessions, the textured track (with reward flavors) was rotated CCW, and distal cues were rotated CW by either 60° (MIS1) or 120° (MIS2), leading to a change in the association between the textured track with reward flavors and distal cues. This experiment creates a reward(flavor)+texture – distal cue mismatch in MIS sessions.

**Figure 7:**
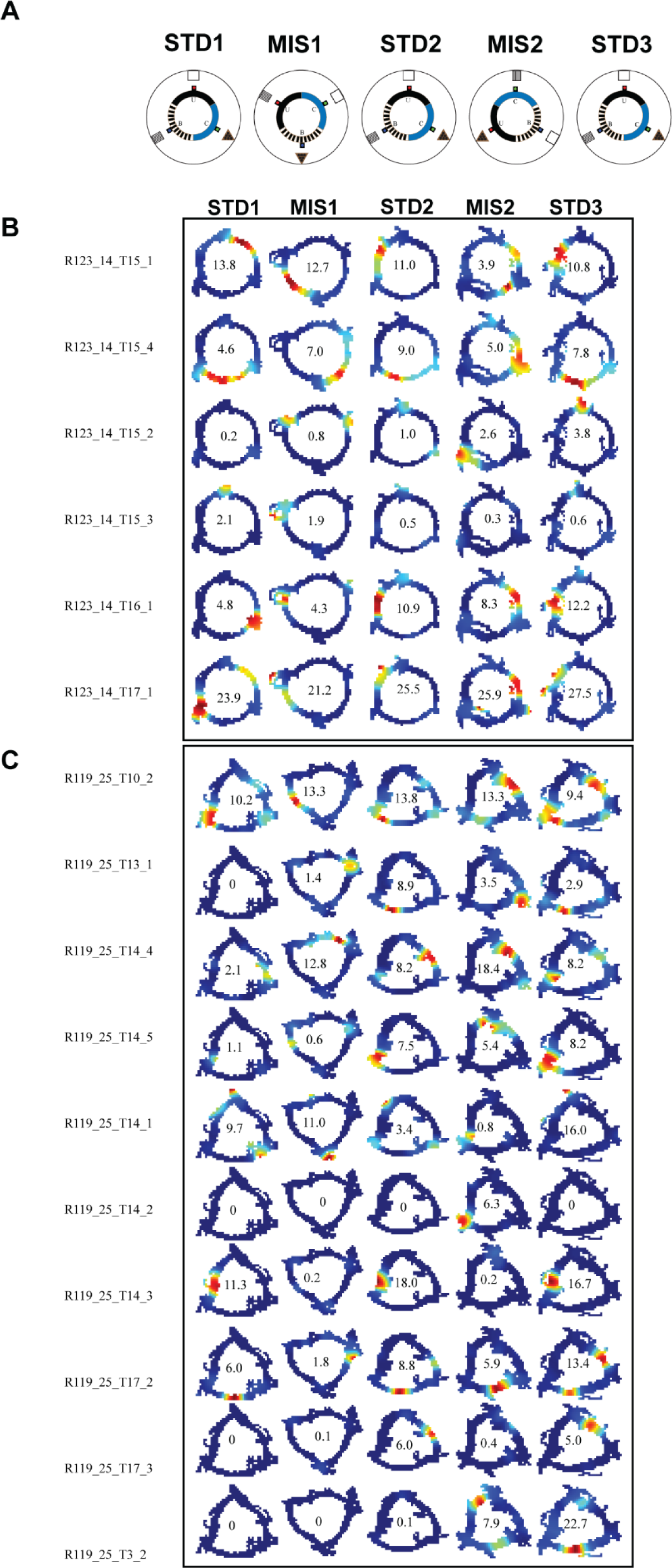
Experimental Paradigm. **A.**Schematic representation of the reward flavor + texture - distal cue double rotation paradigm. Blue dot indicates the spoke where banana flavor pellets(B) were provided, red dot indicates the spoke where Sucrose pellets(U) were provided and green dot indicates the spoke where Chocolate flavored pellets (C) were provided. **B-C** Representative examples of simultaneously recorded CA1 firing rate maps from two ensembles (from different rats) across standard and mismatch sessions. Values inside the rate map indicates the peak firing rate (in Hz). The color coding of the rate map is the same as discussed in the figure legend of Figure 2.

The animal ran CW as in experiment 1, where specific distal cues are aligned to the reward cues and local visual cues. Place cells from two different ensembles were shown in **Fig. 7(B-C)**. In the ensemble shown in **Fig .7B**, some cells maintained stable representation across all three STDs (R123_14_T15_4) where as many cells do not maintain a stable representation (R123_14_T15_1, R123_14_T16_1, R123_14_T17_1). In STD1 vs MIS1 comparison, some cells flipped their firing field at 180° by not orienting with the rotation of the cues (R123_14_T15_1, R123_14_T16_1) where as other place fields rotated CCW following reward flavors (R123_14_T15_4). In STD2 vs MIS2 comparisons, some cells rotated CW following the rotation of distal cues (R123_14_T16_1, R123_14_T17_1) while other cells rotated CCW following the rotation of reward flavors (R123_14_T15_4, R123_14_T15_2). Other cells show weak/no firing at some sessions (R123_14_T15_3), indicating heterogeneous responses of the hippocampal neurons.

**Fig. 7C** shows another ensemble from a different rat. In this ensemble too, hippocampal neurons show diverse responses. Some neurons show stable representation across STDS (R119_25_T10_2, R119_25_T14_3) while other neurons undergo remapping between STDs (R119_25_T17_2). In. MIS1 sessions, we observed the rotation of place field following CW rotation(R119_25_T14_1) and CCW(R119_25_T14_4) rotation simultaneously. In MIS2 sessions, we also observed a similar trend, where cells rotating CW(R119_25_T14_5) and CCW(R119_25_T13_1) too. Also, we observed cells that appeared in only one MIS session(R119_25_T14_2), fired only in STD sessions(R119_25_T14_3), showing a heterogeneous representation of the simultaneously recorded hippocampal neurons. Out of 89 cells, 61 cells fired in all the sessions. Other cells were either appeared or disappeared in some sessions. It was observed that co-recorded place cells did not exhibit a unified response unlike experiment 1. Instead, place cells bind to either local or distal cues simultaneously or it may disappear or appear in MIS sessions.

To quantify the representational characteristics of the hippocampal population, rotational correlation analysis was performed on all pooled neurons that fired in all the sessions of the experiments (**Fig. 8**A). In the STD1 vs. MIS1 comparison, the angle of the mean vector did not shift in response to the rotation of any cues. Also, the peak angles did not show any significant clustering at any rotation of cues or arbitrary rotation, indicated by a low mean vector length (MVL=0.098, p=0.585). In the case of STD2 vs. MIS2 comparison, the angle of the mean vector showed moderate orientation and peak angles showed moderate yet significant clustering at the rotation of reward cues (MVL=0.37, p<0.01).

**Figure 8.**
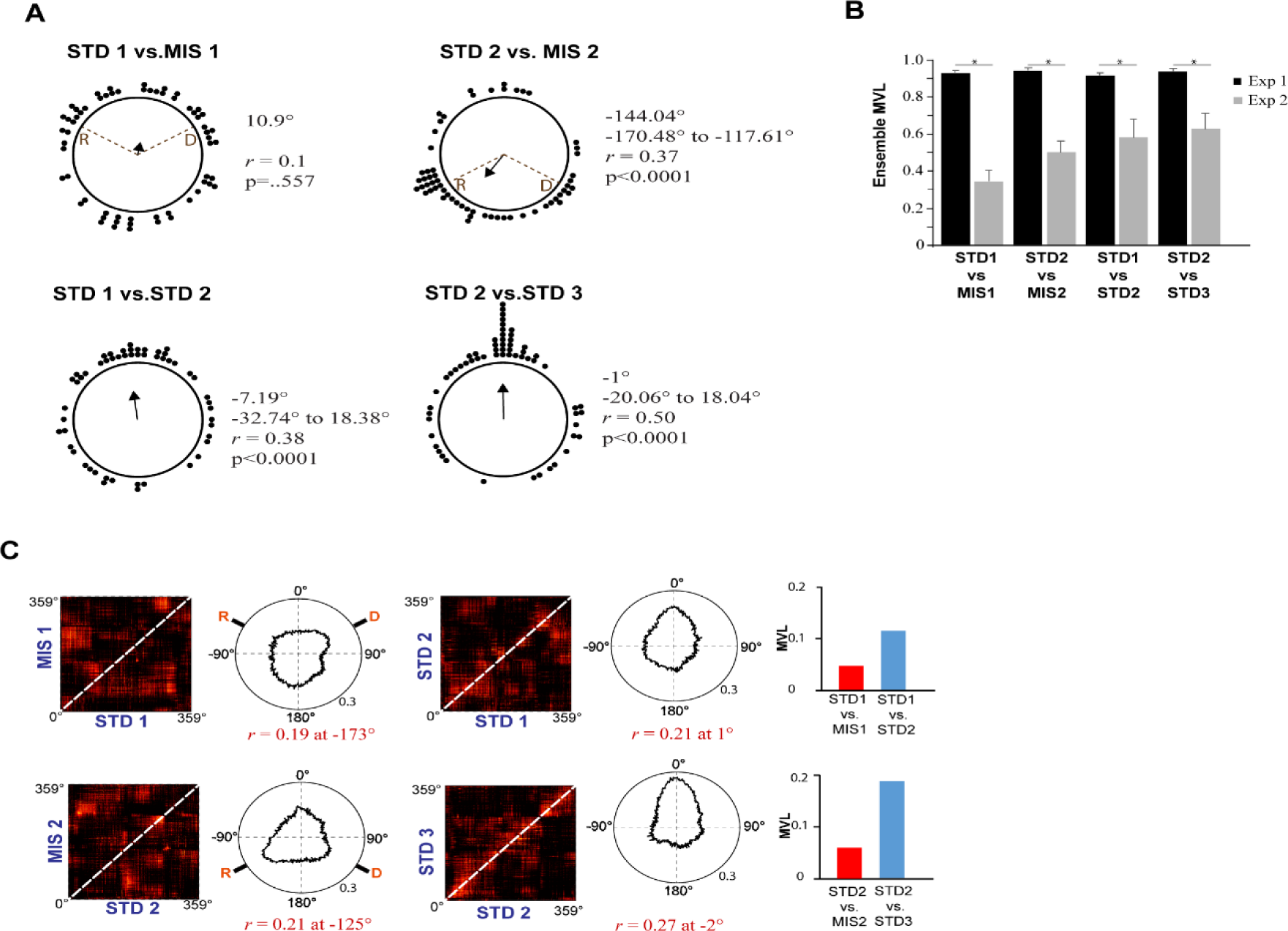
Analysis of individual place field rotation amounts: Comparison with Experiment 1. **A.** Individual rotation of place fields across STD vs MIS and STD vs STD sessions. Brown line indicate the angle of rotation of textured track (and reward flavors) (R) and distal cues (D) in MIS sessions. (each circle represents 1 cell). Values of mean angle of the distribution, 95% confidence interval, length of the mean vector (r) and its significance level (p) is shown near to each plot. In STD1 vs MIS comparison, 95% confidence interval is not shown because p values are not significant. **B.** Comparison of mean vector of all the ensembles in experiment 1 with experiment 2 across STD vs MIS and STD vs STD sessions. * indicates p-value significance (p <0.001, Mann Whitney test). **C.** Correlation matrices from population firing rate vectors of all the cells pooled from each ensemble at each position on the track between STD vs MIS and STD vs STD sessions. The hippocampal activity of STD and MIS comparison is quantified as the polar plot. The values below the polar plot indicates the angle of peak correlation (r) of the hippocampal population. Length of the mean vector (MVL) of each comparison are shown next to the polar plot.

STD vs. STD comparisons showed a similar response where the angle of mean vector was near to 0°. Also, peak correlation angles show significant clustering near to 0° indicating a more stable representation in STD sessions compared to MIS (STD1 vs. STD2: MVL=0.38, STD2 vs. STD3: MVL=0.498, Rayleigh test p<0.01 in both cases). These results showed that, overall, peak correlation angles are dispersed across MIS sessions. This dispersion leads to a low mean vector length of the hippocampal population, suggesting that the hippocampal population exhibited a less coherent, more heterogeneous response in a dynamically changing environment where surface textures are present, which is in line with findings from other studies (Knierim 2002, Lee *et al.,* 2004, Yoganarasimha *et al.,* 2006).

**Fig. 8B** compares ensemble-wise mean vector length in Experiment 1(which is shown in **Fig. 3A**) and Experiment 2 for all STD and MIS comparisons. In Experiment 2, The mean vector length of STD1 vs MIS1, STD2 vs MIS2, STD1 vs STD2 and STD2 vs STD3 comparisons are 0.3375±0.06,0.4944±0.06,0.5744±0.1, and 0.6197±0.09 respectively (mean ± S.E.M.).

Ensemble wise Mean vector lengths of Experiment 2 were significantly lower than Experiment 1(Mann-Whitney test, p<0.001 for all comparisons). This suggests that in Experiment 2, hippocampal representation is less coherent than Experiment 1. To further investigate the location-wise coherency of the hippocampal population, we perform population correlation analysis similar to that of in Experiment 1. Here, population responses between STD vs. STD comparisons and STD vs. MIS comparisons were measured through a 1° bin-wise correlation of linearized firing rate arrays of all the neurons at each position on the track. Population correlation analysis also revealed a lack of specific bands in STD vs. MIS comparisons indicating a heterogeneous population response (**Fig. 8C**). On the other hand, in STD vs. STD comparisons, weak bands were present along the diagonal, indicating that the hippocampal population were more stable across sessions. To quantify population responses across comparisons, a polar plot was constructed from the 2D correlation matrices. In STD1 vs. MIS1 comparison, the angle of peak correlation was not near to any CCW or CW rotation (r=0.19 at -173°). But STD2 vs. MIS2 comparison showed the peak correlation angle near the track rotation, indicating a weak orientation towards the track-based cues (r=0.21 at -125°). STD1 vs. STD2 comparisons showed a peak correlation near zero and a weak band along the diagonal, but the mean vector length is low indicating weak stability (r=0.21 at 1°). In STD2 vs. STD3 comparisons, a band along the central diagonal was present; peak correlation values were near zero with a higher mean vector than other comparisons in experiment 2, indicating a more stable representation (r=0.27 at -2°). But the mean vector lengths in all the comparisons from experiment 2 were lower than the mean vector length from experiment 1 (Mann Whitney test, p=0.0011), suggesting that the hippocampus uses distinct representation in experiment 2.

This observation suggests that the hippocampus follows a heterogeneous representation in the presence of surface textures as shown in the previous studies (Shapiro *et.al*, 1997; Knierim, 2002; Lee *et al*., 2004; Yoganarasimha *et al.,* 2006), even in the presence of reward.

### Attractor dynamics is perturbed in the presence of textures

As we observed a coherent representation in the hippocampus in experiment 1 and heterogeneous representation in experiment 2, we investigated further to compare the network dynamics underlying these representations by studying the attractor properties of the hippocampal system. Specifically, in the case of double rotation experiments, spatial or head directional system is said to exhibit attractor-like properties when the cells in that area maintain spatial offset across sessions, i.e., they are spatially coupled (Knieirim, 2002; Sharma *et al*., 2020). To study the spatial coupling of cells across STD and MIS sessions, we obtained spatial cross-correlation values for each co-recorded place cell pair, and session-wise SXC matrices (rank ordered based on the angle of peak correlation at STD1) were constructed. We compared the spatial offset between co-recorded place cells across five sessions on both experiments. In experiment 1, we observed that SXC peak values were aligned along the diagonal in all the sessions (**Fig. 9A**). This shows that the co-recorded place cells maintain its spatial offset across all the sessions suggesting a robust coupling between place cells. Also, a significant correlation between the mean direction of SXCs of place cells pairs indicates that the hippocampus exhibited attractor dynamics in experiment 1(**Fig. 9B**). In the case of experiment 2, SXC peak values were not aligned along the diagonal in any MIS sessions (**Fig. 9C**). The correlation between the mean direction of SXCs is also significantly very low, showing a lack of coupling due to the perturbance of attractor dynamics in the hippocampus in experiment 2 (**Fig. 9D**).

**Figure 9:**
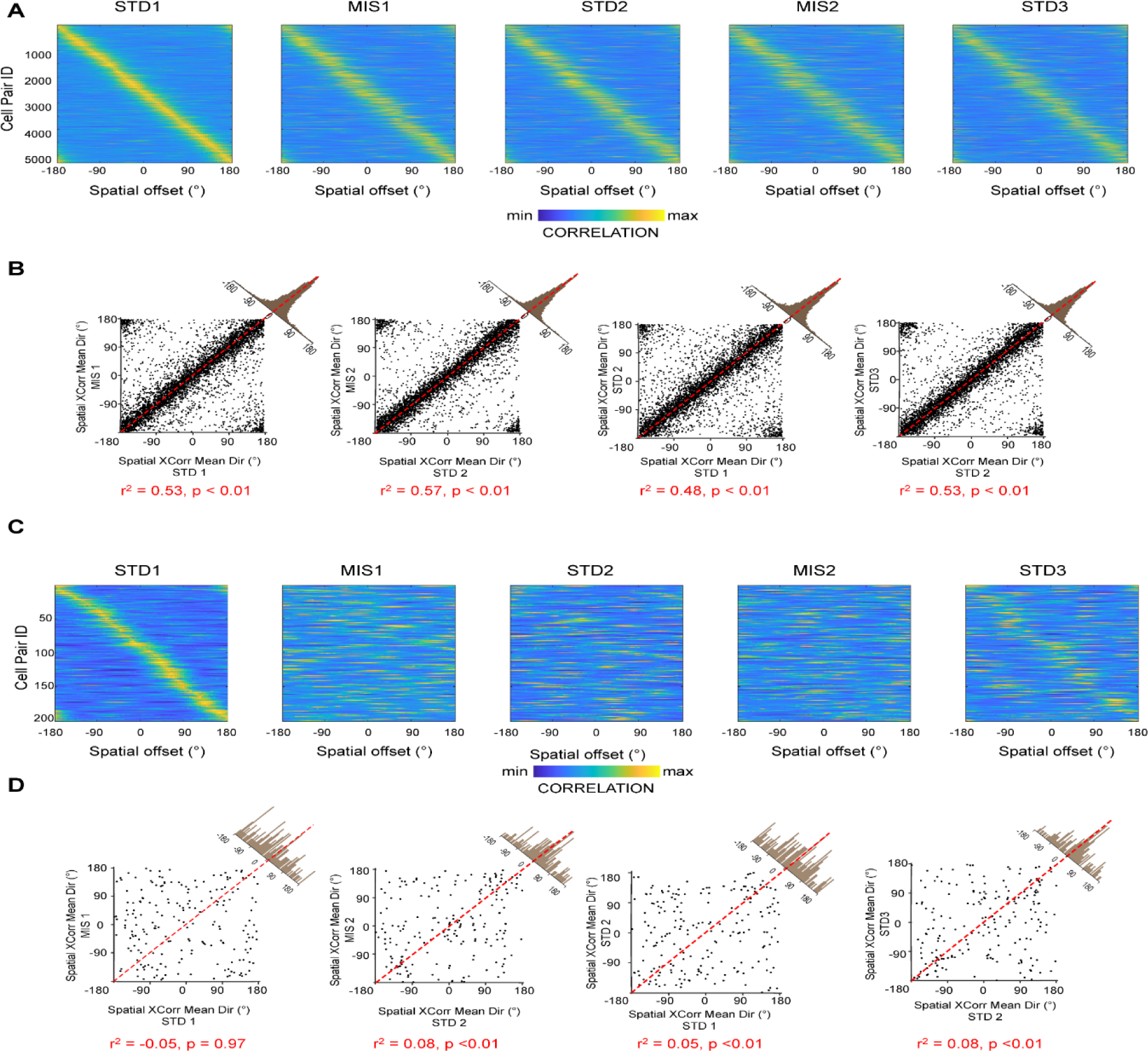
Attractor like spatial coupling in the hippocampus. **A and C**. SXC matrices of co-recorded place cell pairs across three STD and two MIS sessions in experiment 1 (5288 pairs) **(A)** and experiment 2 (201 pairs) **(C)** sorted based on their peak correlation angle values in STD 1 session. **B and D**. Scatter plots showing linear correlation between the mean direction of SXCs of place cell pairs ins STD vs MIS and STD vs STD comparisons in experiment 1(**B**) and experiment 2(**D**). For each of the comparisons, r squared values with its significance level (p) are mentioned below.

These results demonstrate that the hippocampus uses a single coherent attractor-like spatial map in reward(flavor)-distal cue mismatch environments. In contrast, the coherency is reduced in a dynamically changing environment with local visual cues and the hippocampus establishes context-specific spatial maps under these conditions.

## Discussion

This study reports a coherent representation of the hippocampal CA1 ensemble in a reward-distal cue mismatch paradigm. The hippocampal ensembles exhibited a coherent representation even with unstable orientation when the association between the reward flavors and the distal cues is dynamically changed. Across sessions in most experimental days, the place fields dynamically and coherently rotated with the rotation of distal or reward cues. We observed such dynamic responses in most sessions within a day. On some occasions, the ensemble also oriented away from the rotation of distal cues or reward flavors to a fixed angle, whose angular position coincides with the angle of separation between the spokes. The cells were not stable in some STD conditions, yet they coherently reoriented away from 0°. However, in the presence of local visual cues (distinct visual patterns on the track), this coherency is disrupted, leading to the decorrelation of place fields in dynamically changing environments.

Using cue conflict experiments, several studies had reported distinct spatial representation in the hippocampal, parahippocampal, and head direction systems when rats experienced a dynamically changing environment (Knierim & Neunuebel, 2016; Sharma *et al.,* 2020). For example, when distal cues and local cues with distinct visual patterns were rotated to create a varying degree of mismatch in the double rotation paradigm, hippocampal subfields and parahippocampal regions exhibited diverse responses. Medial Entorhinal Cortex (MEC) representations, in general, have followed distal cues and were increasingly decorrelated with increasing local-distal cue mismatch. On the contrary, Lateral Entorhinal Cortex (LEC) representations were weakly controlled by local cues (Neunuebel *et al*., 2013). Both MEC and LEC projects to the Dentate Gyrus (DG) subfield of the hippocampus(Witter,1993). The rate of decorrelation in increasing mismatch angle was more for DG than MEC (Knierim and Neunuebel, 2016). DG produced an enormous change in neural representation with a minor change in EC input; that enables DG to perform as a base for pattern separation. CA3, on the other hand, sustained a coherent spatial representation which gradually degraded as the mismatch angle increased (Neunuebel & Knierim, 2014; Knierim & Neunuebel, 2016). CA3 thus encodes the space to perform pattern completion because of its robust recurrent collateral system (Rolls, 2013). CA3 receives inputs from MEC, LEC, and DG and is controlled by local visual cues. Auto association model of CA3 suggests that one fundamental property of the CA3 recurrent collateral network is its ability to retrieve and complete the whole memory from an input fraction (Rolls & Kesner, 2006; Rolls, 2007). This property enables CA3 in retrieving a coherent representation from the weakly correlated inputs from MEC, LEC, and DG.

Unlike CA3, CA1 lacks strong recurrent collaterals. Therefore, CA1 might integrate information from two distinct pathways (Barrientos and Tiznado, 2016; Knierim and Neunuebel, 2016). One is the direct layer III EC projection to CA1, which is suggested to provide the organism’s current state. The second input (indirectly from EC layer II) to CA1 is from CA3, providing mnemonic representation. The incorporation of both the inputs in CA1 is shown in many experiments as CA1 shows a heterogeneous (incoherent) response where place fields either rotate CW (with distal cues), CCW (with local cues), appear, disappear, remap or even show split representations (Knierim, 2002; Lee *et al*., 2004; Yoganarasimha *et al.,* 2006). This response shows the ability of the CA1 to orient to multiple reference frames simultaneously (Gothard et al., 1996). In addition, CA1 places show more decorrelation than CA3 at smaller mismatch angles(Lee et al., 2004), indicating that CA1 does not reflect as CA3 output. This deviation of CA1 response from CA3 suggests a comparator/mismatch detector function attributed to CA1(Duncan *et al*., 2012) that compares the spatial representation of EC (current state) and mnemonic representation cached in the CA3 (Knierim & Neunuebel, 2016). When different inputs from CA3 and EC reach CA1, these are integrated to exhibit a higher degree of response heterogeneity, encoding specific contexts (Hargreaves *et al*., 2005, 2007).

Hippocampus robustly encodes different goal and reward properties and updates reward information and contingencies as the task progress (Wikenheiser & Redish, 2011; Poucet & Hok, 2017). Studies have reported overrepresentation of place fields at locations near to reward or enhancement of firing near rewards (Hollup *et al*., 2001; Kobayashi *et al*., 2003; Lee *et al*., 2006; Dupret *et al*., 2010; McKenzie *et al*., 2013; Tryon *et al*., 2017). Also, distinct reward-based circuits have been identified that provide context via the CA3-Ventral tegmental area (VTA) pathway and reorganization of reward information via the Locus Coeruleus-CA1 pathway (Luo *et al.,* 2011, Kaufman *et al.,* 2020). The presence of reward cells and conjunctive reward-place cells in the hippocampus suggests the integration reward knowledge onto the hippocampal spatial framework (Gauthier & Tank, 2018; Xiao et al., 2020). Also, studies demonstrated that goal learning distorted entorhinal grid maps by shifting individual grid fields near goal locations (Boccara *et al*., 2019). Additionally, change in navigational strategies for reward restructures the firing rate of MEC cells (Butler *et al*., 2019). Since hippocampal representation is based on the input from MEC and LEC neurons, whenever EC differently encodes distinct environments or contexts, hippocampal neurons should also reflect such change in the spatial representation.

Overall, based on double rotation experiment results and reward processing dynamics of the hippocampus and EC, when the association between reward flavors and distal cues was altered, it is expected that CA1 might create a context-specific spatial map. In contrast, in the experiment 1, when reward flavors and distal cues were rotated to provide different mismatches, we did not observe a heterogeneous response from CA1 ensembles. Instead, CA1 ensembles exhibited a very unstable yet coherent representation, suggesting the interplay of attractor network dynamics. The coherent response might be because of the small change (subthreshold) in EC and CA3 input to CA1 across different mismatch conditions. As the difference in inputs of CA3 and EC might be very small in the mismatch conditions, CA1 might not receive conflicting information. As a result, CA1 could not function as a comparator to differentiate the environments based on contexts. Instead, it may have received information for a similar context. In this scenario, CA1 might have reflected CA3 information arising from the recurrent collateral auto-associative network, featuring coherent representation as a property of the attractor network (Knierim & Zhang, 2012). In the case of experiment 2, when we added local visual cues on the track, the coherency is disrupted, leading to CA1 neurons responding heterogeneously. This response inhomogeneity of CA1 is probably because the addition of visual cues may lead to an increased input change from both EC and CA3 to CA1 across mismatch conditions leading to CA1 neurons differentiating the environments into distinct contexts by showing heterogeneous responses.

As the hippocampal cells is known for encoding location with respect to different spatial reference frames simultaneously (Gothard *et al.,* 1996), one primary reason for a coherent response in experiment 1 might be the absence of continuous local cues. Here, local cues (reward flavors) are provided only at the spokes. Thus, the discretization of the local cue experience may not differentiate the EC and CA3 representations in cue-conflict sessions leading to the coherent representation of CA1, the output region of EC and CA3. Also, as the animal receive pellets on every spoke, the task may not require the animal to encode a change in context.

But the dynamical and coherent switch in the orientation of place fields within an experimental day shows that the hippocampus dynamically binds to different cues. While place cells are known for their stability, the dynamically unstable and coherent place fields reported here suggest a possibility of a shift in the animal’s attention to different reference frames in different sessions because of reward flavors that add salience (and value) parameters in the paradigm. This switch in attention may lead to dynamic binding to multiple reference frames, yet coherently in an experimental day (Muzzio, 2018). It is also shown that hippocampal representation of contexts is modulated by task demands (Zinyuk *et al*., 1999).

As the hippocampus produces distinct spatial maps based on different contexts, a single map in this scenario can be considered a map generalization mechanism, leading to categorizing similar contexts. Recent work by Kinsky and group have shown that place field maps show random and coherent rotation in an exploration task without orienting to specific arena cues (Kinsky *et al*., 2017). Our work also indicates dynamic coherent rotation, but here the task is more goal-directed having specific flavors of rewards, requiring the animal to run in a particular direction for food rewards. Also, our results show that the clustering of angles is not at any random angle but are ∼120° apart, showing cue/spoke based orientations which demonstrate that specificity of spatial reorientation may be guided by track geometry (Keinath *et al*., 2017).

The addition of local visual cues enabled the CA1 to differentiate it as different contexts in dynamically changing environments. Recent studies have shown that the hippocampus place cells encode the boundaries of local surface textures (Wang *et al*.,2020) and spatial coding resolution was increased in the presence of local visual cues (Bourboulou *et al*., 2019).

Additional studies also point out the rotation of a subset of place fields with local cues and the appearance and disappearance of place fields in dynamically changing environments (Shapiro *et al.,* 1997; Knierim, 2002; Lee *et al.,* 2004). These studies show the critical role of textures (local visual cues) in differentiating contexts.

In the first experiment, place cell ensembles, on many occasions, have oriented with the rotation of reward flavors. But the presence of reward flavors alone could not create a context-specific spatial map because either it is not a continuous cue like local visual cues (which is present everywhere on the track) or it may not have a strong impact on pattern separation like local visual cues. The breakaway from a robust attractor to a heterogeneous representation in the second experiment also suggests the importance of visual cues in categorizing spatial contextual memory.

One potential shortcoming of the present study is that we did not record CA3 and DG assemblies together to study their representation. As DG is shown to perform pattern separation and CA3 as pattern completion, it will be interesting to study how the hippocampal representation and orientation changes in cur-conflict associated with reward flavors.

## Author Contributions

I.R.N. conceived and designed the study, performed the experiments and data analysis. I.R.N. and D.R interpreted the data. I.R.N. and D.R. wrote the manuscript.

## Data and Code Accessibility

The data that supports the findings and the codes used for data analysis are available from the corresponding author upon request.

## Acknowledgements

The authors thank Apoorv Sharma for constructive feedback towards proofreading the manuscript, and Shiladitya Laskar for assisting during behavioral experiments.

---

**Supplementary Figure 1-1.**
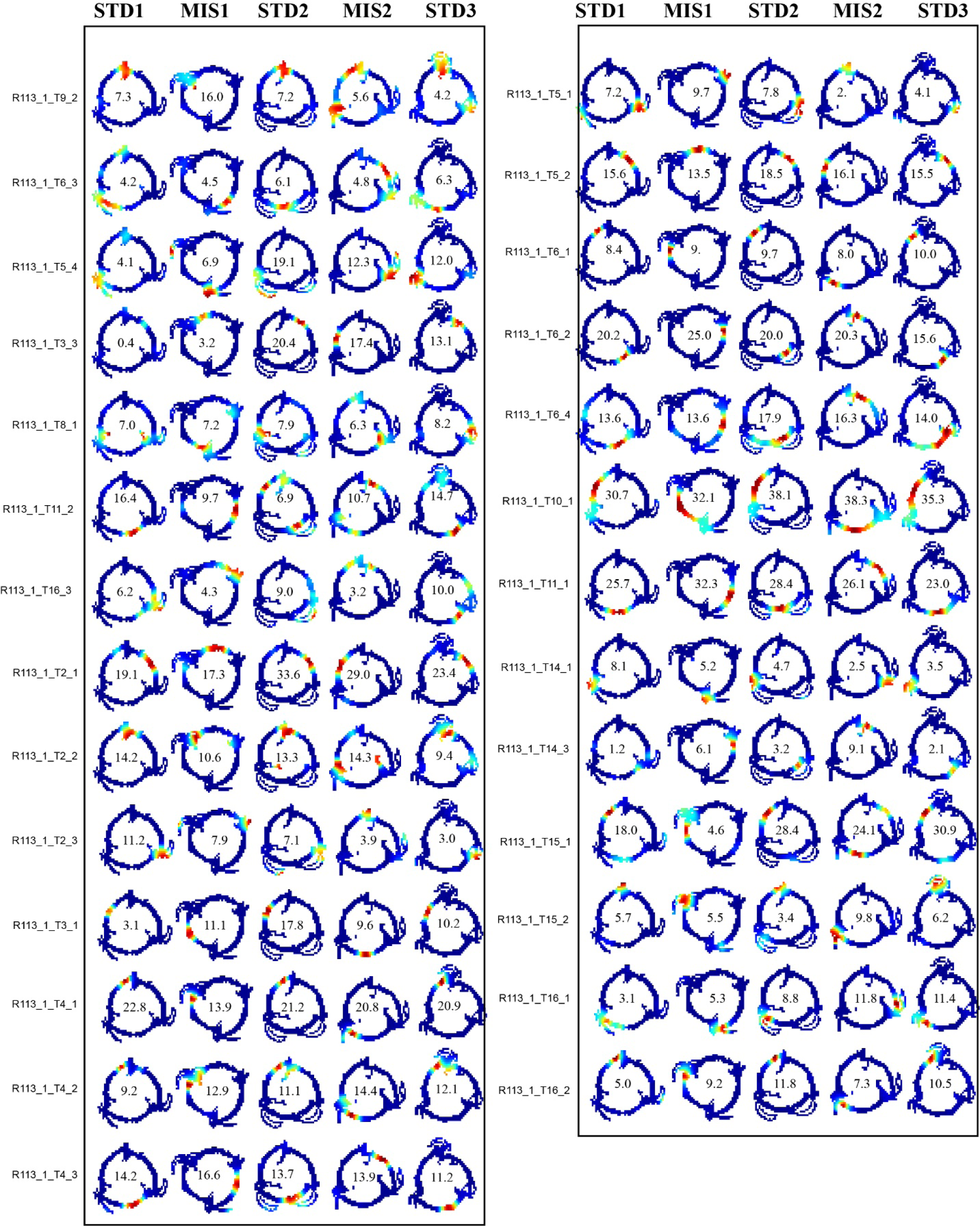
Rate maps of all the neurons present in ensemble-1 across STD and MIS sessions from Rat-113. Left side of the plot shows the IDs of each neurons. For example, in R113_1_T9_2 ID: R113 indicates Rat-113, 1 indicates experimental day, T9 indicates tetrode 9 and 2 indicates the cluster number from that tetrode.

**Supplementary Figure 1-2.**
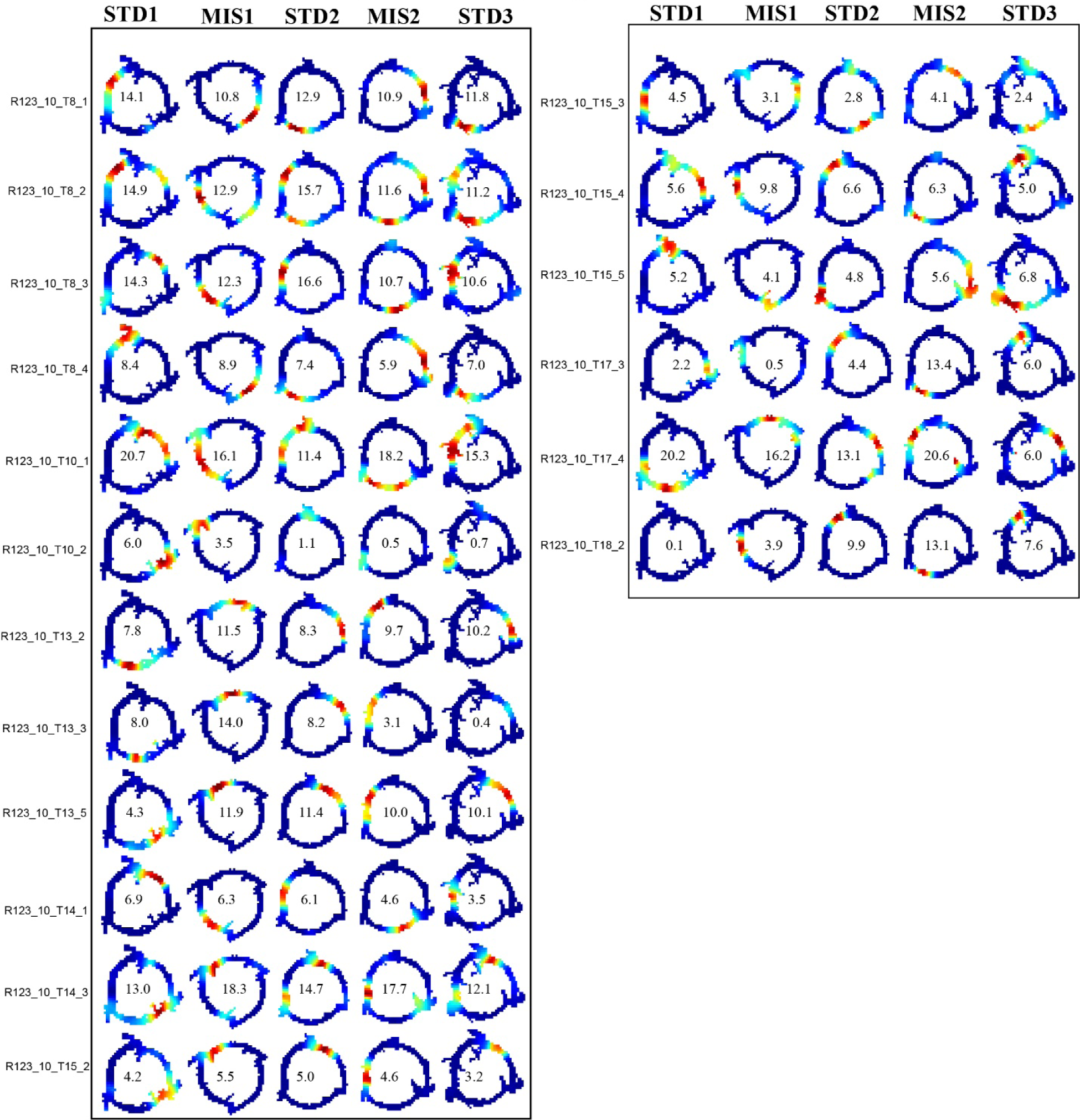
Rate maps of all the neurons present in ensemble-3 across STD and MIS sessions from Rat-123. Left side of the plot indicates the IDs of each neurons.

